# DeepCNA: an explainable deep learning method for cancer diagnosis and cancer-specific patterns of copy number aberrations

**DOI:** 10.1101/2024.03.08.584160

**Authors:** Mohamed Ali al-Badri, William CH Cross, Chris P Barnes

## Abstract

Chromosomal instability leads to an increased rate of chromosome or part-chromosome copy gains or losses. Referred to as chromosome copy aberrations (CNAs), these mutations are highly prevalent in cancer cells and contribute to abnormal genome structure, genetic diversity within a tumour, drug resistance, and evolution to metastatic disease. Within each cancer type, it remains unclear how CNAs define each particular malignant phenotype, yet some key events have been characterized. Here, we present a novel deep learning method, DeepCNA, and apply it as part of an exploration of CNAs present across 7,500 whole genomes and 13 cancer types. DeepCNA is an explainable AI approach, that reveals both established and novel loci that contribute to cancer type. It can be used as a diagnostic tool in situations such as cancer of unknown primary site, and it can discover CNAs that are prognostic. It also has several applications for researchers, including drug target and biomarker discovery.

## Introduction

Chromosomal instability (CIN) is a phenomenon observed in most cancer cells and it is defined as an increased rate of changes in both the number and structure of chromosomes. This form of genomic instability results from a wide range of mechanisms, including those directly associated with the mitotic spindle apparatus and chromosome mis-segregation^1^, but also less well understood events such as dysfunction of telomeres^2^, impairments in cell cycle checkpoints^3^, deficits in double stand break repair^4^, and genome doubling^5^. Chromosome copy aberrations (CNAs), the specific loci that result from CIN, have been observed in around 90% of solid cancer genomes^6^ making them one of the most prominent features of the cancer genome. The observed chromosome profile of a given cancer is a product of these mechanisms plus its unique evolutionary walk, which is influenced by positive selection forces from the microenvironment (defined in a major part by the site of origin) along with the stochastic nature of mutagenesis. For this reason, the specific combination of CNAs found in cancer subtypes has been studied in detail.

Following the historical work of von Hansemann and Theadore Boveri^7–9^ and more recently *in vitro*^10^, *in vivo*^11^, and bioinformatic studies^12^ a view of CIN as a general potentiate of malignancy has been constructed^13^. More recent studies have sought to explore the genome-wide architecture that results from CIN^14^ and to explain patterns of CNAs through mutational signatures^15,16^. In the case of point mutations, signatures are thought to underlie the differing mutagenic processes^17^, and while this concept is appealing in the analysis of CNAs there are many hurdles, including the definition of data features and how these might represent the underlying processes of mutagenesis.

One of the challenges associated with CIN is its complexity. A core aim for researchers is to classify those sites that are important to cancer biology, separating from those that are passengers and therefore resultant from stochastic forces. An ideal situation is that specific sets of CNAs define each cancer subtype and correlate with patient outcome, thus acting as biomarkers. Such events may also yield valuable gene targets for drug developers. To date, many cancer-specific CNAs have been identified^18,19^ and in specific gene contexts, such as *TERT* amplifications^20^ they can act as biomarkers for disease severity, and in other cases drug resistance^21^, or as novel drug targets^22^. That said, these CNAs are often discovered from frequency or mechanistic studies and it remains challenging to survey sufficient numbers of whole cancer genomes and invoke appropriate methods to discover loci that are critical to each specific cancer type and subtype.

In this study we have undertaken a new approach whereby we examined CNA patterns using the latest AI methodologies. Using DeepCNA, an explainable AI, we classified cancers based exclusively on CNA patterns then dissected the loci that contributed information to the classification. Our hypothesis is that certain CNAs capture differences between cancers and contribute to the varying biological context of differing cancer subtypes. We also test the prognostic information of our findings using patient outcome data, thus demonstrating how the method could be employed in a variety of research settings.

## Results

### Analysis of 7522 cancer genomes from the 100,000 Genomes Project

The 100,000 Genomes project was managed by Genomics England and completed in 2019^23,24^. It served as a transformative project for integrating whole genome sequencing (WGS) into the National Health Service (NHS). Participants provided consent for their genomic data to be associated with annonymised longitudinal health records, and the subsequent data was made available to researchers via a secure virtual environment. Clinical value as been demonstrated from pan-cancer analyses that investigated the distribution of genome-level drivers and made correlations with outcome^25^, though a detailed analysis of CNAs has yet to be performed.

The dataset contains 13,000 cancer-normal paired genomes with linked outcome data, and each genome is labelled with a diagnosis. Diagnosis labels that represent a mixture of biological systems (for instance sarcoma) were omitted from this study, since it is unlikely that they will be classifiable using CNAs under a single label. Furthermore, rare cancer subtypes with fewer than 100 genomes (for instance endocrine) were also removed since the class signal would likely be undetectable when analysed alongside the more frequent cancer subtypes. Since CNAs are the only input to DeepCNA, we also excluding those genomes with less than 30% of the genome altered, as quantified by the Percentage Genome Altered (PGA). This threshold aligns with previous work suggesting that a substantial genomic alteration is a reliable indicator of increased chromosomal instability^26,27^. Finally, we excluded genomes that had undergone a very high degree of rearrangement, since these are likely to involve distinct biological mechanisms such as chromothripsis^28^ or chromoplexy^29^ which are out of scope. A high degree of chromosome rearrangement would also interfere with functioning of the neural network underlying DeepCNA, since this was based on individual chromosome inputs (as outlined below). This resulted in 7522 genomes for analyses (Table 1).

**Table 1:**
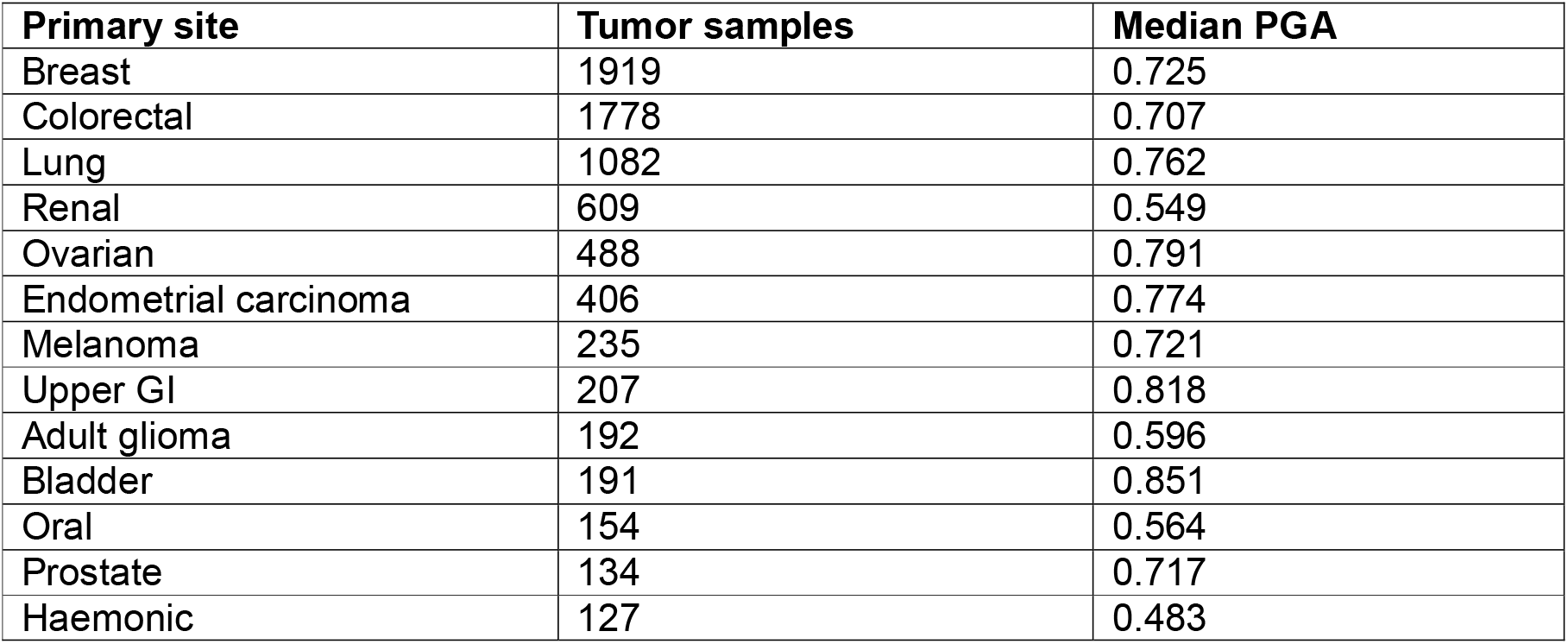
Distribution of tumour types in the GEL data sets together with their median percentage genome altered (PGA).

### A novel deep learning framework for analysis of cancer genomes

We trained the neural network underlying DeepCNA which takes CNA data as input and classifies the genomes based on primary site label. To prepare the data, we split the genomes into 100kb bins and calculated the median total copy number, plus the median minor copy state within each bin. The minor copy state is not phased but is assigned by the majority of CNA calling algorithms for loci where there is a deviation in allele frequency in the overlapping single nucleotide changes. Together this reduced each genome into a 28749 × 2 tensor and provides a conserved genome coordinate across all genomes. The data were then scaled feature-wise across samples such that each bin had zero mean and unit variance. The neural network architecture was designed as an ensemble of 22 chromosome-specific sub-networks, separately acting on each chromosome in the input (Figure 1). As such, each chromosome sub-network has an input layer of nodes equal to the binned size of the respective chromosome, with chromosome 1 the largest number of nodes and chromosome 21 with the fewest. These chromosome feature extractors then serve as inputs to a deep feed-forward network. Each genome in our dataset is composed of a large feature set, while the classification involves the 13 classes, represented by each diagnosis label. Navigating through such a large feature space demands a robust model capable of identifying relevant patterns while avoiding overfitting. Therefore, we implemented a hyperparameter optimisation approach using 5-fold cross validation (see Methods). The chromosome feature extraction prevented overfitting, as did dropout layers with high probability (p = 0.4). In addition, we chose not to use convolutional layers to improve downstream interpretability.

**Figure 1:**
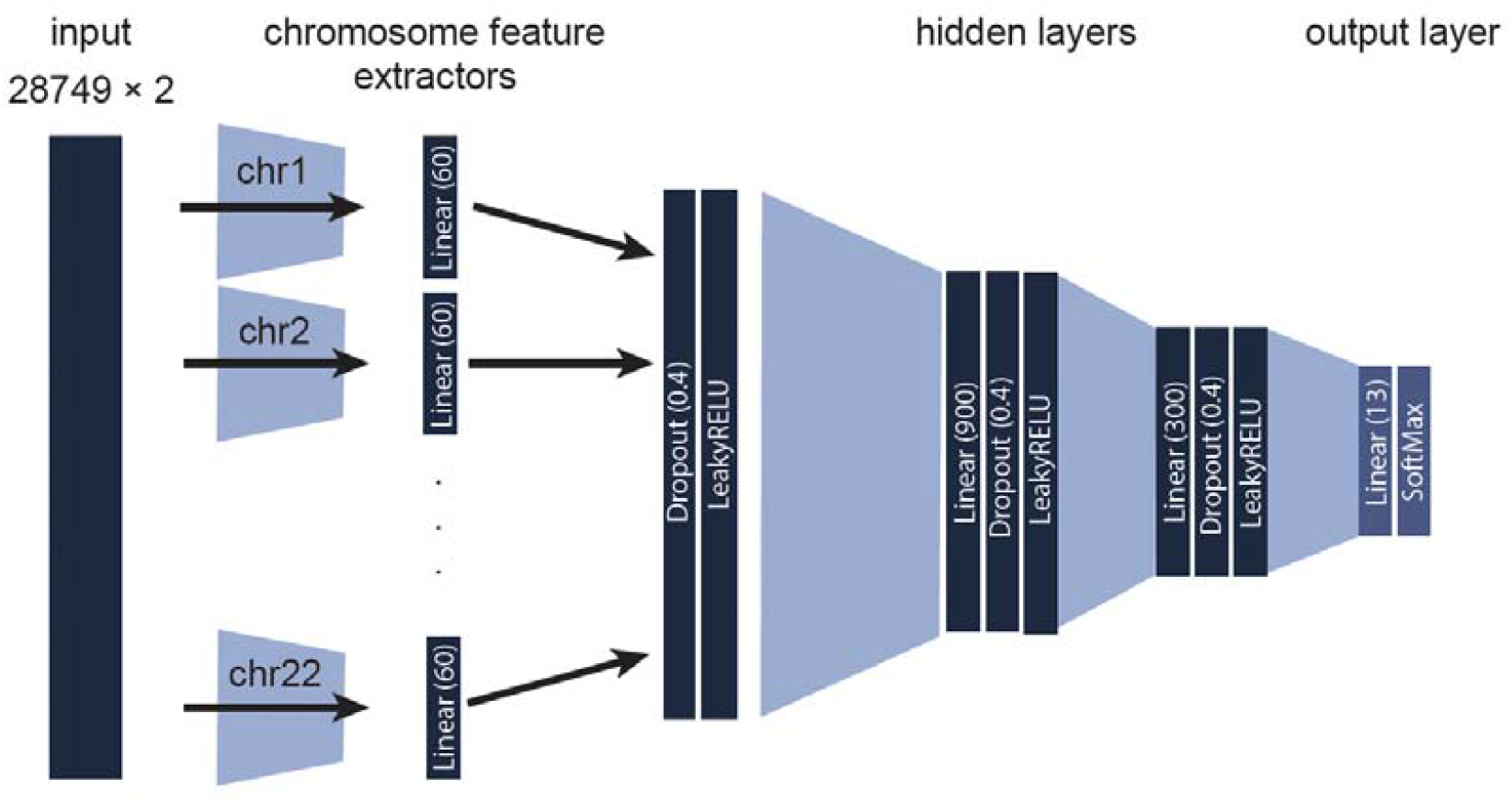
Schematic of the neural network architecture, illustrating the input from whole genome sequencing (WGS) samples. Each chromosome’s binned data feeds into a sub-network, culminating in a deep feed-forward network with hierarchical layers.

### Cancer diagnoses can be determined from chromosome number aberrations

DeepCNA was able to classify cancers based exclusively on CNA data, with average accuracy of 70% across the entire dataset (Figure 2A). As expected, the accuracy was variable across cancer type, ranging from 85% in colorectal cancer (CRC) to 54% in bladder. This pattern in performance can largely be explained by sample size (Figure 2B). We also explored the performance as a function of median PGA (Figure 3C). This showed a peak at intermediate PGA values suggesting that some level of variation is required to distinguish genomes, but too much reduces the discriminatory power. The cancer types with the highest median PGA are bladder, upper gastrointestinal and ovarian. Despite these general trends, they are not universally applicable to all cancers. For example, lung cancer has a relatively low classification accuracy (71%) despite having a large sample size. This could be due to the heterogeneity of lung cancer within the data. Interesting properties of model confusion occur also, which may be due to the overlap in primary sites and therefore biology with other cancers. For example, endometrial carcinoma and ovarian cancer show high misclassification rates with each other (16% and 13%). Similarly, we observed some misclassification between CRC and upper gastrointestinal cancers (misclassification rate of 11%), which also have a degree of biological overlap. These results show the performance of DeepCNA in classifying genomes based on the CNA distribution and that there are cancer specific differences in chromosomal copy number patterns.

**Figure 2:**
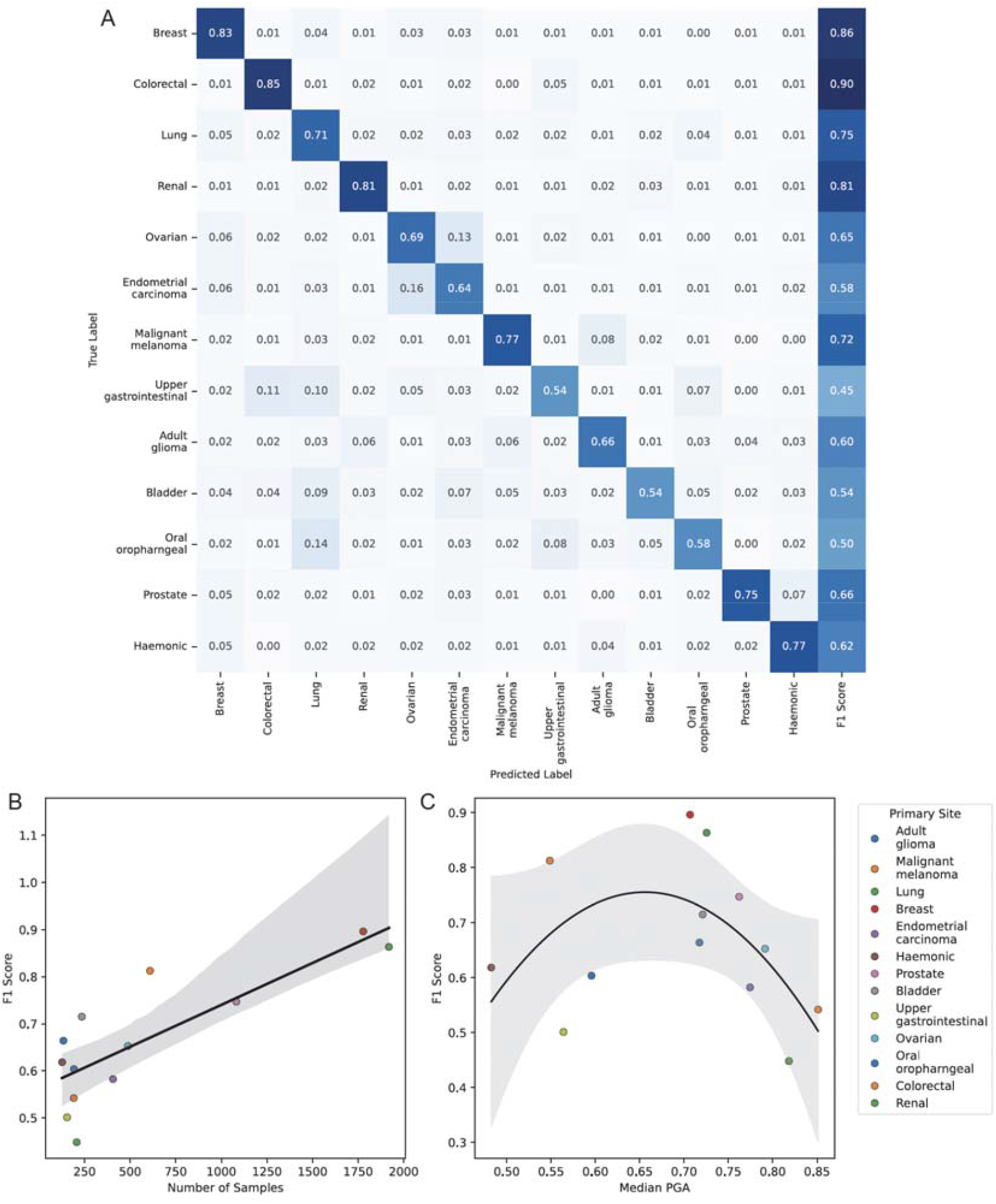
Performance of the fitted model. A) Confusion matrix for each cancer primary site, accompanied by F1 score, providing a detailed assessment of the model’s classification performance. B) F1 score against sample size with a linear regression line. C) F1 score against median PGA with a second-order polynomial regression line. Shaded regions in B,C indicates 95% confidence. F1 scores and the confusion matrix are averaged over test sets from 5-fold cross validation.

**Figure 3:**
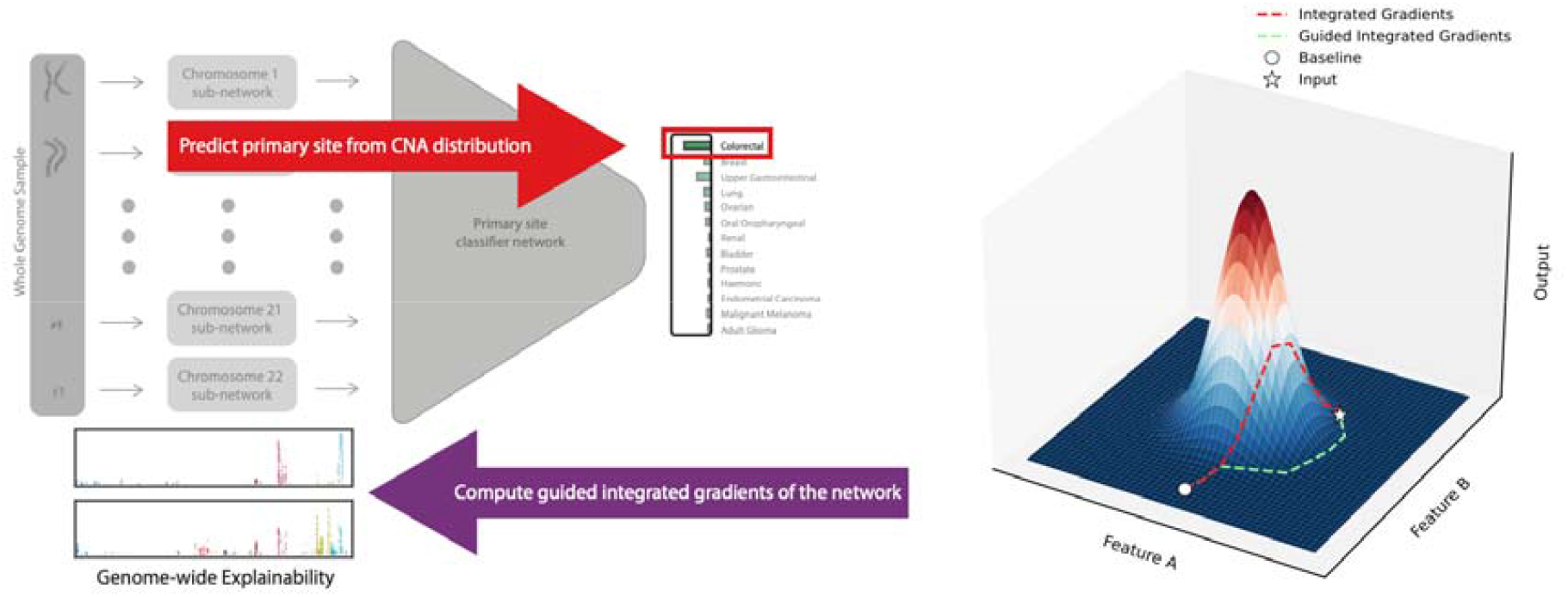
Depiction of the Guided Integrated Gradients (GIG) method, tracing the process from sample classification prediction to genome-wide explainability. A comparison is drawn between Integrated Gradients (IG) and GIG, highlighting the path selection mechanism in GIG for enhanced gradient accumulation analysis.

### Determining functional variation across the whole genome

In order to infer the most important CNAs that determine diagnosis, we employed an approach based on guided integrated gradients (GIG)^30^, which is a technique used in explainable artificial intelligence to understand the predictions of neural networks by attributing importance scores to input features^31^ (Figure 3, Methods). By utilising GIG, our aim was to provide a more nuanced understanding of the features contributing to the classification of primary sites, enhancing the interpretability of the model’s decision-making process. The GIG inference using the trained model was applied individually for each genome, generating a ranking of genomic bins. DeepCNA was trained exclusively on the CNA states of each bin, with none of the associated metadata such as gene or enhancer content fed into the training. By aligning the inferred feature importance with specific genomic annotations and examining whether a feature corresponds to a specific CNA (e.g. duplication, deletion, LOH), we were able to define biological context, and obtain an understanding of the model’s decision-making. In principle this could be used in future investigations to capture distinctions between any set of cancers, including those that represent genetic, histological, or prognostic subtypes.

### Recovery of CNAs notable to cancer evolution in individuals

Sample-level explainability may reflect those CNAs that contribute to cancer evolution. Accordingly, we reviewed the CNA distributions across each individual patient and examined those with high attribution scores in either the total or minor copy features. Representative genomes from patients diagnosed with either breast, lung, and CRC are shown below, and a complete summary is available in the supplementary materials (also see Methods). The 10 highest CNAs for each sample and their corresponding copy state, cytoband, COSMIC oncogenes and enhancers are shown.

DeepCNA correctly reported that genome 4885 was a Breast cancer with a classification probability of 0.96 (Figure 4A). In this genome several CNAs, represented as distinct but sometimes proximal loci, were discovered. Some CNAs have been previously reported in Breast cancer or other cancers, and included amplifications at 8p11.22^32,33^ and 16p13.12, the latter being associated with *ERCC4* and novel in the context of breast cancer. The 19q12 locus was also amplified and is the location of nine genes including cyclin E1 (*CCNE1*), a potential cancer driver^34^. The method also reported several CNAs across the 17q arm, which is known to contain *AXIN2*^35,36^ and loci previously linked to poor prognosis^37^.

**Figure 4:**
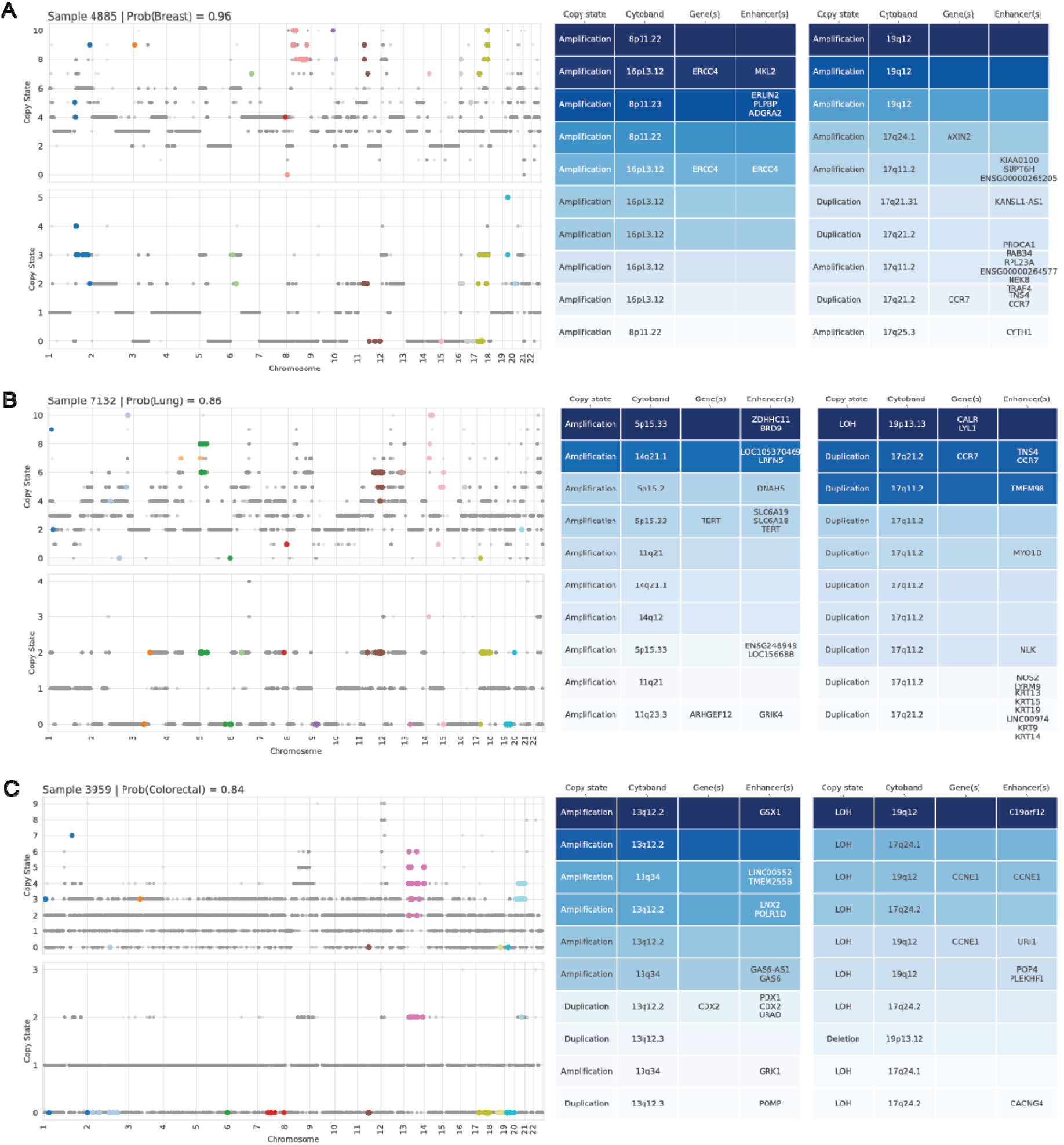
AI explainability in single whole genome sequencing samples across different cancer types. (A) One breast cancer sample, (B) one lung cancer sample, and (C) one colorectal cancer sample, each showcasing CNAs across the genome. The top part of each panel displays the total copy number for both alleles, while the bottom shows the minor allele copy number. High-importance genomic loci, identified by the model, are colour-coded by chromosome, with unselected loci in grey. Accompanying tables detail the copy state (Diploid, Duplication, Amplification, Deletion, LOH), cytoband, presence of oncogenes, and enhancers for the high attribute loci in both total and minor allele signals. Table rows are colour-coded based on the magnitude of normalised attribution values from the GIG analysis, highlighting key loci contributing to the AI’s classification decision.

In another example, genome 7132 was classified correctly as a Lung cancer with a probability of 0.86 (Figure 4B). The highest attribution scores included chromosomes 5, 11, 14, 17, along with LOH CNA in chromosome 19. In particular, the amplification at 5p15.33 encompasses the genes *TERT* and *CLPTM1L*, implicating their potential role in lung cancer biology^38–40^ along with 5p12.2 which has been studied in non-small cell lung cancer. CNAs identified on the 14q13.2–14q21.1 bands are interesting since these linked to the *MBIP, NKX2-1, NKX2-8*, and *PAX9*, genes which have been implicated in lung cancers in never smokers^41^. The association of 17q11.2 with *HER2*, an important biomarker in breast cancer^42–44^.

Finally, in genome 3959, our model correctly suggested the CRC diagnosis with a probability of 0.84 (Figure 4C). The amplification of 13q12.2, specifically involving the *CDX2* gene, corroborates a previous study that implicates this gene in the Wnt/β-catenin signalling pathway, which is critical to CRC biology^45^. Likewise, on the 13q34 cytoband *IRS2* has previously been identified as an oncogene, offering a potential mechanism for PIK3 kinase pathway activation in CRC^46^. As with the Breast cancer sample, our model suggested that the 17q24.1 region, and the association of *AXIN2*^47^ was potentially important in this particular instance of CRC, demonstrating the value that certain CNAs have across several cancer types, but also within specific patients.

### Cohort level analysis reveals chromosome arms implicated in specific cancer diagnoses

As well as the ability to recover known and novel CNAs within individuals, we explored the high attribute CNAs across the entire cohort, first focusing on whole chromosome arms, by aggregating all ranked CNAs based on their frequency (see Methods). A subset of outputs from DeepCNA from the breast, lung and CRC cancer cohorts are shown below (Figure 5A-C). To visualise the attribution distributions across all samples within a cohort and depict the arrangement of consistently highly ranked features, we also show stacked Manhatten plots and heatmaps (Supplementary Figures).

**Figure 5:**
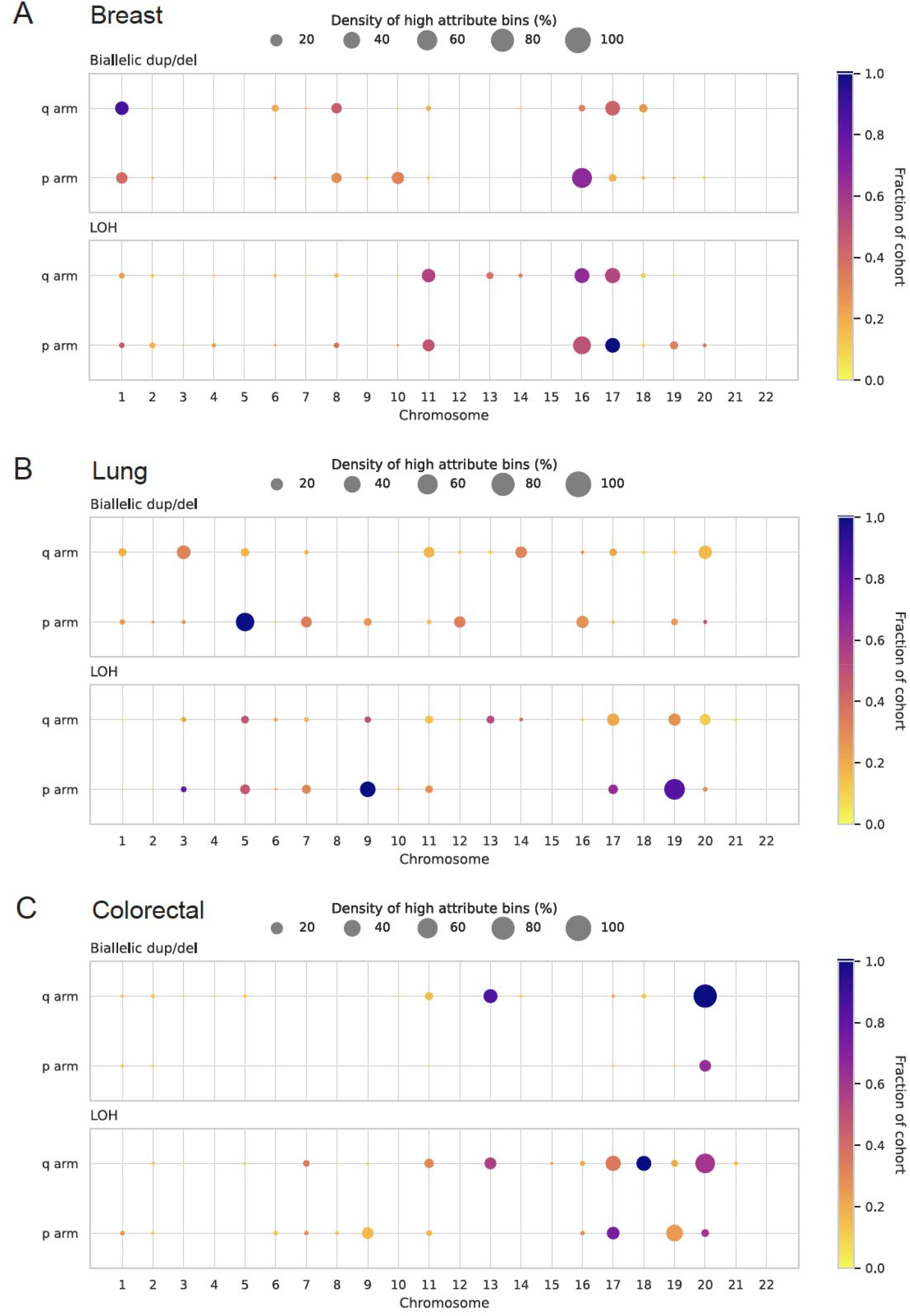
Chromosome arm level analysis of genomic alterations and AI-driven attribution in cancer cohorts. (A-C) Bubble plots for breast, lung, and colorectal cancer cohorts for both total and minor allele copy numbers, displaying high-attribute loci across chromosome arms. Bubble size represents the density of significant bins, with colour indicating the fraction of the cohort sharing these loci.

To explore these results in detail, we extracted the top ranked cytobands for breast, lung, and CRC cohorts by selecting all events present in > 50% of samples, resulting in 18, 10 and 28 loci respectively. We used literature searches to find established links with these regions (see Supplementary Tables 1-3). Here we provide a very brief summary to demonstrate how researchers can use this method to identify chromosome arms of interest across specific cancer types. Together such findings demonstrate how potentially novel biological details, but also established CNAs, can be found through machine learning principles, that are completely naïve to the biological underpinnings.

In breast cancer, regions 1q, 16p, and 17q exhibited the highest density of attribution scores, with the latter being a known initiation event in breast and other cancer types^48^ (Figure 5A). Interestingly, 1q has been reported as a translocation partner of 16p^49^ and these copy events may be suggestive of a further biological link. Moreover, the 11q loci was implicated in this dataset, and deletions have been association with relapse in patients with lymph node– negative breast cancer who did not receive anthracycline-based chemotherapy^50^.

In lung cancer, 5p, 9p and 19p arms were among the highest density (Figure 5B). Gains on 5p have been identified as a frequent even in small cell lung cancer (SCLC)^51–53^, while 9p has also been highlighting in lung cancer pathogenesis^54^. Simultaneous mutations at *LKB1* and *BRG1* are common in lung cancer cells, and it may be that 19p is highlighted in our analysis due to its link with heterozygosity (LOH) and specific tumor suppressor genes such as these^55^.

In CRC we found high attribute scores at 13q, 17p, 18q, and 20q (Figure 5C). These events have been consistently observed in large-scale genomic studies, and each loci may pertain to a variety of biological effects. For example, gain of 20q is observed in more than 65% of CRCs and harbours multiple genes of interest, such as *BCL2L1, AURKA*, and *TPX2*^56–59^. Additionally, *PLAGL2* and *POFUT1* expression patterns have been linked copy number amplification on 20q^57,60,61^. Likewise *SMAD4*, present in 18q21.1, has a well-established link with CRC pathogenesis^62–65^.

### Diagnosis prediction scores act as prognostic markers

Another opportunity in DeepCNA is the discovery of new prognostic markers based on the two primary outputs: the high attribute score CNAs and the cancer prediction scores. We hypothesise is that CNAs discovered within a particular cancer type have higher attribution scores when they induce phenotypes associated strongly with a particular disease. Although only limited numbers of CNAs currently have prognostic value, it is possible that an AI approach recovers information that have been missed by conventional studies. We also considered that the prediction scores reflect the genomic closeness of a particular cancer to an archetype, or optimum karyotype for invasive cellular behaviour, which may therefore also have prognostic significance.

To test these hypotheses, we performed an outcome analysis using standard Kaplan Meier and Cox Proportional Hazard (CPH) statistics, restricting each test set to a given cancer type and removing Stage 1 diseases which are routinely curable (see Methods, n = 5168). Including a stage correction for the remaining cases, we found that 42 loci significantly affected patient outcome, though following a strict Bonferroni-based multiple testing correction, this was reduced to 4 loci (6q23.1, 6q23.2, 10p14, 13q34) all in breast cancer (Figure 6A). We noted that the hazard ratio of these loci was only slightly lower than stage in the CPH model (stage: HR = 2.2, 6q23.1: HR = 2.0, Figure 7A), demonstrating the value that CNAs discovered via DeepCNA have in some diagnostic settings.

**Figure 6:**
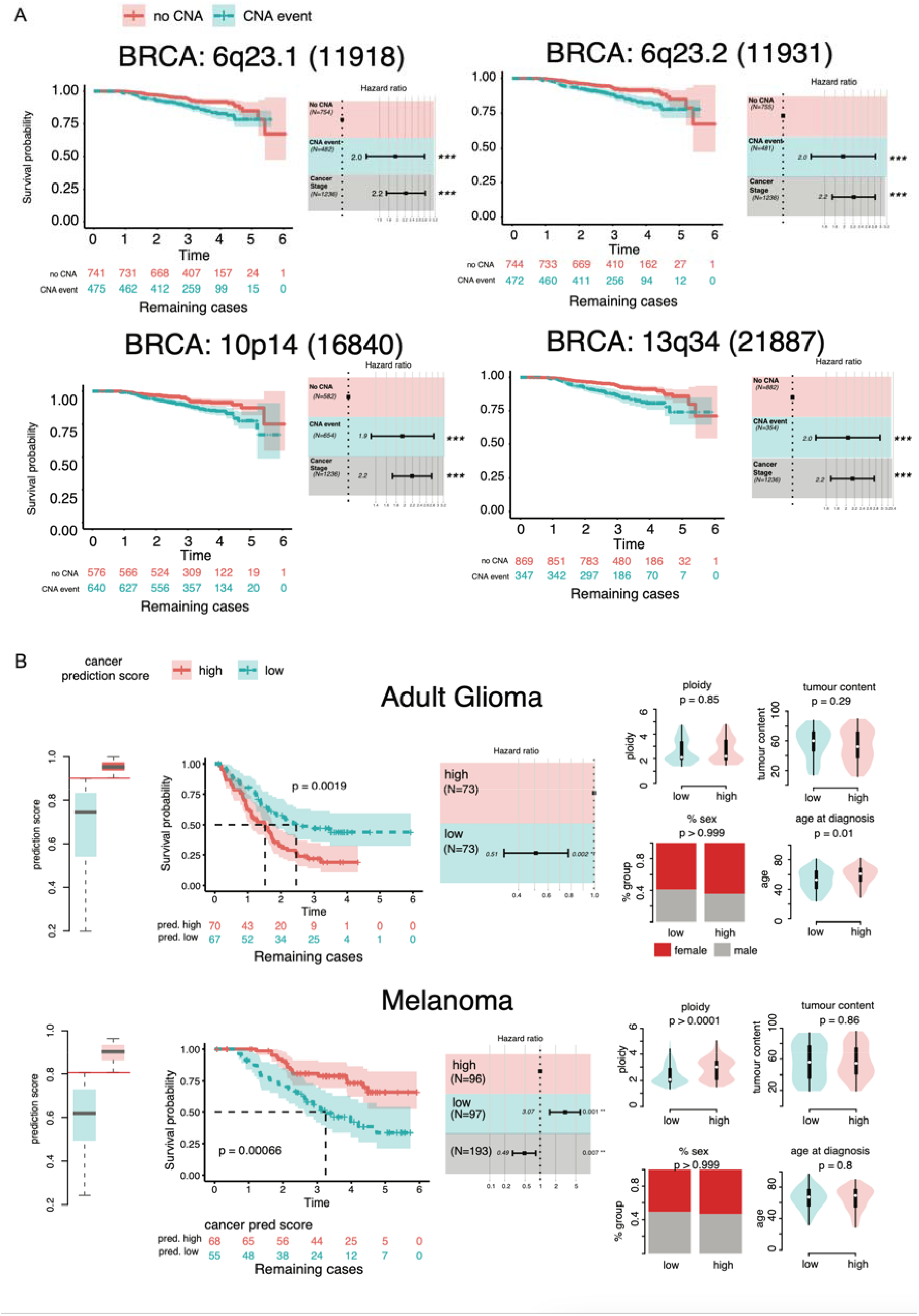
Analysis of patient outcome A) 42 loci with significant scores effect patient outcome following stage correction. Four loci remained significant following multi-testing correction (chromosome location and bin ID in brackets labelled). Kapalan Meier plots and tables with patient numbers shown on left, with forest plots stage corrected on the right. B) Splitting of patient groups was performed using the median cancer scores (left most plots). Kaplan Meier plots show significant impact on outcome in adult glioma and melanoma (note that glioma does not use a staging system thus absent from forest plots). Right most plots show other genetic features of the patient cohorts, including ploidy, tumour content of the biopsy, proportion of male or female patients, and age at diagnosis.

The cancer prediction scores generated by DeepCNA also have prognostic value in higher stage diseases in some cancer types. By partitioned each set of patients into high and low cancer prediction scores using the median across each cancer type (Figure 7B), we found that a high prediction score pertained to a worse prognosis in adult glioma, when compared to low scores (Figure 6B). Interestingly, we also found that a low prediction score pertained to a worse prognosis in melanoma (Figure 6B). We wondered whether this link between prediction score and outcome might be reflecting some underlying clinical or genetic factors, and in examining ploidy, tumour content, age, and sex, across these patients, we found that the high-low Glioma groupings were different by age (p = 0.01). In melanomas a significant distinguishing feature between the high-low cancer prediction score was ploidy (p = 0.0001). This highlights the complex relationships between the different features of cancer evolution and differences in patient outcome, and how AI methods can help to reveal novel metrics and biomarkers.

## Discussion

Although deep learning approaches have previously been used to classify cancer genomes^66^, and to explore cancers of unknown primary^67^, none have explored CNAs and none have been applied to differing cancer types on this scale. In this study we developed DeepCNA, an explainable, unbiased AI that can be used to diagnose specific cancer types with a high degree of accuracy, based on CNA information alone. It can also be used as part of a research strategy to discover novel biomarkers and biological features related to specific CNAs, and it can reveal both individual patient and cohort-wide information. It also has potential to be used as part molecular diagnostics and the identification of cancer-specific drug targets.

As means of demonstration we applied DeepCNA to 7500 genomes from the 100,000 Genomes Project. We showed that cancers can be classified from patterns of CNA alone, and that there are interesting biological relationships between some cancer types that have an anatomical relationship, such as ovarian and endometrial cancers. We consider that one application of DeepCNA could be to diagnose cancers of unknown primary, which represent a clear clinical conundrum from a therapeutic and treatment perspective^67^. It could also be used to better understand complex cancers that encompass a large number of subtypes with unknown or disputed biological overlap, such as sarcomas^68^.

DeepCNA aligns with broader efforts to enhance personalised and effective treatment strategies in oncology, marking a significant contribution to the field of genomic medicine. That said, there are some limitations associated with explainable AI approaches, including the fact that the usefulness of the information depends on the predictive performance of the model on that sample^69^. Future work could involve the use of unsupervised learning approaches that can alleviate these issues.

In summary, the effective diagnosis of cancer is possible from CNA data alone, which is a testament to their significance in cancer biology. DeepCNA serves as a valuable tool for further research on CNAs, potentially revealing novel aspects of cancer biology important for medical intervention.

## Methods

### Data and pre-processing

The Genomics England 100,000 Genomes Project was an NHS transformation project that took place between 2016 and 2019^70^. During this time between 19,000 and 20,000 participants with cancer consented for the study and had paired blood and tumour biopsies DNA sequenced to 50 and 100X respectively. The bioinformatic processing utilised an accredited bioinformatic pipeline, which included industry-standard tools and in house quality^71^.

The Genomics England (GEL) CNA data using in this project was produced using the CANVAS tool^72^. We visualised the relationship between the median proportion of the genome impacted by LOH and the median number of LOH segments in each cancer primary site by mapping genomic states (such as deletion, diploid, LOH, duplication and amplification) to integers. This map-ping facilitated the identification and quantification of continuous stretches of each copy state within the samples. LOH segments were specifically extracted and analysed to determine their distribution across the genome. The degree of whole genome doubling (WGD) was calculated using the PCAWG approach^73^ GEL primary sites with fewer than 100 sample are insufficient for training a neural network (testicular, endocrine, etc.) or those whose class labels is composed of multiple primary sites or does not fall under the same convention (Hepatobiliary pancreatic, sarcoma, cancer of unknown primary site, etc.) were omitted from the analysis. Furthermore, we determined the threshold for CNA-driven cancers to have at least 30% of their genome altered (PGA). The CNA calls for each patient were organised into tabulated data indicating the CNA state by genomic coordinate for both alleles, along with the major allele. To prepare these data for use by the neural network, we conducted a series of transformations. Each sample was anonymised and processed to generate an individual array, wherein the copy state of the 22 autosomes was binned into 100 kb segments, mapped to a single array element. Within each sample, two channels were encoded: one for the total allele CNA state and another for the minor allele CNA state. The resulting array has shape (2, 28749). This data structure has a conserved genome coordinate across all samples, where each element/bin denotes the copy state of a given genomic region of 100 kb. Prior to training the network, the dataset was feature-wise normalised by subtracting the mean and dividing by the standard deviation.

### Deep learning approach

In order to maximise interpretability, we chose not use convolution operations. The network architecture was composed of an initial ensemble of 22 chromosome-specific sub-networks, separately acting on each chromosome in the input (Figure 2). As such, each chromosome sub-network has an input layer of nodes equal to the binned size of the respective chromosome and an output number of nodes, denoted *n*_*chrom*_. The chromosome feature extractors then serve as inputs to a deep feed-forward classifier network. This classifier network comprises of three layers with input given by *22n*_*chrom*_, two hidden layers of size *n*_*h*1_, *n*_*h*2_ respectively. Each layer is composed of linear, dropout, and LeakyReLU layers. The output layer, with 13 nodes, aligns with the number of cancer primary sites in the GEL dataset and employs a softmax function to produce the final predictions.

During training, a weighted sampler was used to address the class imbalance in the GEL data (see Table 1). The neural network was trained over 50 epochs using the stochastic gradient descent optimiser with cross-entropy loss function. Learning rate, momentum, batch size, dropout rate and network structure (*n*_*chrom*_, *n*_*h*1_, *n*_*h*2_) were determined in a hyperparameter tuning process using the Optuna framework^74^. This used 5-fold stratified cross validation to evaluate the model performance for each parameter combination. The precise ranges of hyperparameters and their selected values are shown in Table 2. All performance metrics such as F1-scores and confusion matrices were calculated over the 5-folds to reduce bias. In a final step, and for downstream explainability, the model was fit with optimal parameter combination to the whole data set. The neural network was trained using PyTorch^75^.

**Table 2:**
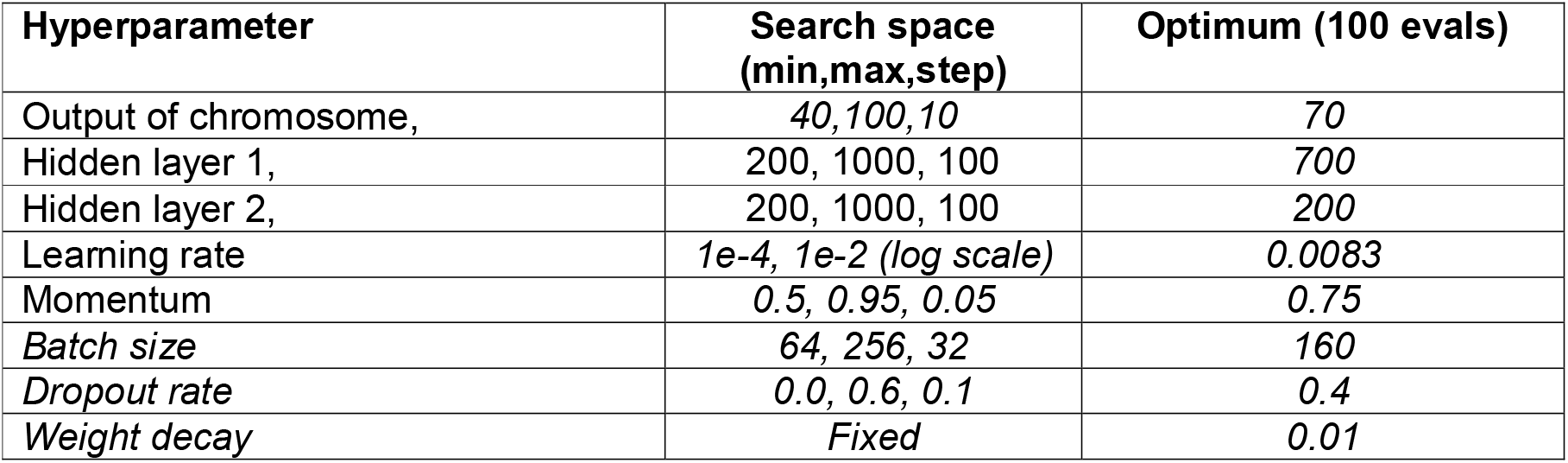
Hyperparameter optimisation results.

### AI explainability

In order to infer the most important loci that determine diagnosis, we employed an approach based on Integrated Gradients (IG). IG is a technique used in explainable artificial intelligence (xAI) to understand the predictions of neural networks by attributing importance scores to input features^31^. Although there have been a number of improvements to the approach^76^, it can still be challenging to interpret the information gained. We used the guided integrated gradients (GIG) method^30^, since unlike conventional IG, adaptive path methods reduce the noise accumulated along the IG path by defining a path integral between the baseline and input sample. Conditioning the path not just on the input sample but also on the model being explained. This conditioning ensures a more accurate attribution of feature importance, considering the specific characteristics of the neural network’s decision boundaries. The resulting feature attributions are numerical scores, ranking the input features based on their influence on the model’s prediction. This ranking gives the relative importance of each genomic locus in contributing to the classification of an individual cancer primary site versus the other primary sites. In our analysis, the input data is highly disparate across samples, therefore the explainability results can be sensitive to the choice of baseline. Namely, a conventional choice of baseline for image data may take the form of a black image or an array of zero values, which is counterproductive for data composed of (feature-normalised) CNA states. Instead, the median CNA state of a sample across a given channel is chosen as a baseline to start the GIG path integral, ending with the sample CNA state values.

### Cohort-level explainability analysis

In order to aggregate the explainability score across each primary site cohort we selected the top 1% of loci within a sample and then selected those resulting loci that were present in more than 5% of the cohort. This conservative approach allowed us to visualise patterns across the primary sites and to verify known biological features (Figure 6). To provide a more statistically robust assessment of the significant loci for the clinical outcome analysis, we performed an association study following previous work^76^. This comprised a loci dependent t-test on the raw attribute signal where the null distribution was a sample of attribute scores at that loci in other primary sites. The null distribution contained a number of points equal to the number of samples in the test cohort. To reduce variability in this testing procedure we repeated the t-test ten times and calculated the average p-value. The p-values were corrected using the Bonferonni multiple testing correction. Loci were further processed to remove chromosome arm level regions leaving only focal events for the clinical outcome analysis.

### Clinical outcome analysis

For the majority of patients’ outcome data was available, including date of death and time interval of followup. We focused on higher stage disease which are associated with fatal conditions (stage *>* 1, n = 5168). Data from the high attribute loci and cancer prediction scores were subjected to Kaplan-Meier (log rank) and cox proportional hazard testing, the latter correcting for can-cer stage in all type bar Glioma. The R Survival and Survminer packages were used for this purpose, and multiple testing correction was performed using the Benjamin Hochberg method on any reported p-values. Visualisation was handled by the standard plotting functions included in the packages: ggsurvplot, and ggforest^77^.

## Data availability

The data used for all the analysis, together with summaries of the outputs are available in a Zenodo repository: https://zenodo.org/records/14892622

## Code availability

The code used to train and optimise the models, perform the guided integrated gradients analysis and make all the plots are available in the following GitHub repository: https://github.com/ucl-cssb/DeepCNA

## Acknowledgments

CPB and MAB acknowledge funding from the Wellcome Trust (209409/Z/17/Z). The authors acknowledge the use of the UCL Myriad High Throughput Computing Facility (Myriad@UCL) and the UCL Department of Computer Science High Performance Computing Cluster, and associated support services, in the completion of this work.

## Notes

### Competing Interest Statement

The authors have declared no competing interest.

### Summary of Updates

Main text updated to include more detailed methods; Figures updated

## References

1. Hosea, R., Hillary, S., Naqvi, S., Wu, S. & Kasim, V. The two sides of chromosomal instability: drivers and brakes in cancer. Signal Transduct Target Ther 9, 75 (2024).

2. Murnane, J. P. Telomere dysfunction and chromosome instability. Mutat Res 730, 28–36 (2012).

3. Simonetti, G., Bruno, S., Padella, A., Tenti, E. & Martinelli, G. Aneuploidy: Cancer strength or vulnerability? Int J Cancer 144, 8–25 (2019).

4. Lieber, M. R. The mechanism of double-strand DNA break repair by the nonhomologous DNA end-joining pathway. Annu Rev Biochem 79, 181–211 (2010).

5. Sanz-Gómez, N., González-Álvarez, M., De Las Rivas, J. & de Cárcer, G. Whole-Genome Doubling as a source of cancer: how, when, where, and why? Front Cell Dev Biol 11, 1209136 (2023).

6. Weaver, B. A. & Cleveland, D. W. The aneuploidy paradox in cell growth and tumorigenesis. Cancer Cell 14, 431–433 (2008).

7. Boveri, T. Concerning the origin of malignant tumours by Theodor Boveri. Translated and annotated by Henry Harris. J Cell Sci 121 Suppl 1, 1–84 (2008).

8. Holland, A. J. & Cleveland, D. W. Boveri revisited: chromosomal instability, aneuploidy and tumorigenesis. Nat Rev Mol Cell Biol 10, 478–487 (2009).

9. Wright, N. A. Boveri at 100: cancer evolution, from preneoplasia to malignancy. J Pathol 234, 146–151 (2014).

10. Lengauer, C., Kinzler, K. W. & Vogelstein, B. Genetic instabilities in human cancers. Nature 396, 643–649 (1998).

11. Rayi, A. & Hozayen, S. Chromosome Instability Syndromes. in StatPearls (StatPearls Publishing, Treasure Island (FL), 2025).

12. Dewhurst, S. M. et al. Tolerance of whole-genome doubling propagates chromosomal instability and accelerates cancer genome evolution. Cancer Discov 4, 175–185 (2014).

13. Baker, T. M., Waise, S., Tarabichi, M. & Van Loo, P. Aneuploidy and complex genomic rearrangements in cancer evolution. Nat Cancer 5, 228–239 (2024).

14. Watkins, T. B. K. et al. Pervasive chromosomal instability and karyotype order in tumour evolution. Nature 587, 126–132 (2020).

15. Steele, C. D. et al. Signatures of copy number alterations in human cancer. Nature 606, 984–991 (2022).

16. Drews, R. M. et al. A pan-cancer compendium of chromosomal instability. Nature 606, 976–983 (2022).

17. Steele, C. D., Pillay, N. & Alexandrov, L. B. An overview of mutational and copy number signatures in human cancer. J Pathol 257, 454–465 (2022).

18. Goh, J. Y. et al. Chromosome 1q21.3 amplification is a trackable biomarker and actionable target for breast cancer recurrence. Nat Med 23, 1319–1330 (2017).

19. Qiu, X. et al. MYC drives aggressive prostate cancer by disrupting transcriptional pause release at androgen receptor targets. Nat Commun 13, 2559 (2022).

20. Hugdahl, E., Kalvenes, M. B., Mannelqvist, M., Ladstein, R. G. & Akslen, L. A. Prognostic impact and concordance of TERT promoter mutation and protein expression in matched primary and metastatic cutaneous melanoma. Br J Cancer 118, 98–105 (2018).

21. Leonetti, A. et al. Resistance mechanisms to osimertinib in EGFR-mutated non-small cell lung cancer. Br J Cancer 121, 725–737 (2019).

22. Pons, G. et al. Analysis of Cancer Genomic Amplifications Identifies Druggable Collateral Dependencies within the Amplicon. Cancers (Basel) 15, 1636 (2023).

23. Turnbull, C. et al. The 100L000 Genomes Project: bringing whole genome sequencing to the NHS. BMJ 361, k1687 (2018).

24. 100,000 Genomes Project Pilot Investigators et al. 100,000 Genomes Pilot on Rare-Disease Diagnosis in Health Care-Preliminary Report. N Engl J Med 385, 1868–1880 (2021).

25. Sosinsky, A. et al. Insights for precision oncology from the integration of genomic and clinical data of 13,880 tumors from the 100,000 Genomes Cancer Programme. Nat Med 30, 279–289 (2024).

26. Thu, K. L. et al. Lung adenocarcinoma of never smokers and smokers harbor differential regions of genetic alteration and exhibit different levels of genomic instability. PLoS One 7, e33003 (2012).

27. Pikor, L., Thu, K., Vucic, E. & Lam, W. The detection and implication of genome instability in cancer. Cancer Metastasis Rev 32, 341–352 (2013).

28. Kloosterman, W. P. et al. Chromothripsis is a common mechanism driving genomic rearrangements in primary and metastatic colorectal cancer. Genome Biol 12, R103 (2011).

29. Shen, M. M. Chromoplexy: a new category of complex rearrangements in the cancer genome. Cancer Cell 23, 567–569 (2013).

30. Kapishnikov, A. et al. Guided Integrated Gradients: An Adaptive Path Method for Removing Noise. Preprint at 10.48550/arXiv.2106.09788 (2021).

31. Sundararajan, M., Taly, A. & Yan, Q. Axiomatic Attribution for Deep Networks. Preprint at 10.48550/arXiv.1703.01365 (2017).

32. Adélaïde, J. et al. Chromosome region 8p11-p21: refined mapping and molecular alterations in breast cancer. Genes Chromosomes Cancer 22, 186–199 (1998).

33. Voutsadakis, I. A. 8p11.23 Amplification in Breast Cancer: Molecular Characteristics, Prognosis and Targeted Therapy. J Clin Med 9, 3079 (2020).

34. Natrajan, R. et al. Functional characterization of the 19q12 amplicon in grade III breast cancers. Breast Cancer Res 14, R53 (2012).

35. Xu, X., Zhang, M., Xu, F. & Jiang, S. Wnt signaling in breast cancer: biological mechanisms, challenges and opportunities. Mol Cancer 19, 165 (2020).

36. Zeng, Y. A. & Nusse, R. Wnt proteins are self-renewal factors for mammary stem cells and promote their long-term expansion in culture. Cell Stem Cell 6, 568–577 (2010).

37. Latham, C. et al. Frequent co-amplification of two different regions on 17q in aneuploid breast carcinomas. Cancer Genet Cytogenet 127, 16–23 (2001).

38. McKay, J. D. et al. Lung cancer susceptibility locus at 5p15.33. Nat Genet 40, 1404– 1406 (2008).

39. Cq, Z. et al. Amplification of telomerase (hTERT) gene is a poor prognostic marker in non-small-cell lung cancer. British journal of cancer 94, (2006).

40. Ju, K., Sh, K., Kc, K., Jw, P. & Jm, K. Gain at chromosomal region 5p15.33, containing TERT, is the most frequent genetic event in early stages of non-small cell lung cancer. Cancer genetics and cytogenetics 182, (2008).

41. B, J. et al. Genomic aberrations in lung adenocarcinoma in never smokers. PloS one 5, (2010).

42. Rakha, E. A. et al. Updated UK Recommendations for HER2 assessment in breast cancer. J Clin Pathol 68, 93–99 (2015).

43. Biserni, G. B., Engstrøm, M. J. & Bofin, A. M. HER2 gene copy number and breast cancer-specific survival. Histopathology 69, 871–879 (2016).

44. Kallioniemi, O. P. et al. Association of c-erbB-2 protein over-expression with high rate of cell proliferation, increased risk of visceral metastasis and poor long-term survival in breast cancer. Int J Cancer 49, 650–655 (1991).

45. Salari, K. et al. CDX2 is an amplified lineage-survival oncogene in colorectal cancer. Proc Natl Acad Sci U S A 109, E3196–3205 (2012).

46. Day, E. et al. IRS2 is a candidate driver oncogene on 13q34 in colorectal cancer. Int J Exp Pathol 94, 203–211 (2013).

47. Otero, L. et al. Variations in AXIN2 predict risk and prognosis of colorectal cancer. BDJ Open 5, 13 (2019).

48. Reinholz, M. M. et al. Breast cancer and aneusomy 17: implications for carcinogenesis and therapeutic response. Lancet Oncol 10, 267–277 (2009).

49. Rye, I. H. et al. Quantitative multigene FISH on breast carcinomas identifies der(1;16)(q10;p10) as an early event in luminal A tumors. Genes Chromosomes Cancer 54, 235–248 (2015).

50. Girish, V. et al. Oncogene-like addiction to aneuploidy in human cancers. Science 381, eadg4521 (2023).

51. Ju, K. et al. High frequency of genetic alterations in non-small cell lung cancer detected by multi-target fluorescence in situ hybridization. Journal of Korean medical science 22 Suppl, (2007).

52. Schwendel, A. et al. Primary small-cell lung carcinomas and their metastases are characterized by a recurrent pattern of genetic alterations. Int J Cancer 74, 86–93 (1997).

53. Romeo, M. S. et al. Chromosomal abnormalities in non-small cell lung carcinomas and in bronchial epithelia of high-risk smokers detected by multi-target interphase fluorescence in situ hybridization. J Mol Diagn 5, 103–112 (2003).

54. Sato, M. et al. Identification of chromosome arm 9p as the most frequent target of homozygous deletions in lung cancer. Genes Chromosomes Cancer 44, 405–414 (2005).

55. Rodriguez-Nieto, S. & Sanchez-Cespedes, M. BRG1 and LKB1: tales of two tumor suppressor genes on chromosome 19p and lung cancer. Carcinogenesis 30, 547–554 (2009).

56. Carvalho, B. et al. Multiple putative oncogenes at the chromosome 20q amplicon contribute to colorectal adenoma to carcinoma progression. Gut 58, 79–89 (2009).

57. Li, D. et al. PLAGL2 and POFUT1 are regulated by an evolutionarily conserved bidirectional promoter and are collaboratively involved in colorectal cancer by maintaining stemness. EBioMedicine 45, 124–138 (2019).

58. Sillars-Hardebol, A. H. et al. TPX2 and AURKA promote 20q amplicon-driven colorectal adenoma to carcinoma progression. Gut 61, 1568–1575 (2012).

59. Ali Hassan, N. Z. et al. Integrated analysis of copy number variation and genome-wide expression profiling in colorectal cancer tissues. PLoS One 9, e92553 (2014).

60. Du, Y. et al. POFUT1 promotes colorectal cancer development through the activation of Notch1 signaling. Cell Death Dis 9, 995 (2018).

61. Su, C. et al. Studying the mechanism of PLAGL2 overexpression and its carcinogenic characteristics based on 3’-untranslated region in colorectal cancer. Int J Oncol 52, 1479–1490 (2018).

62. Alazzouzi, H. et al. SMAD4 as a prognostic marker in colorectal cancer. Clin Cancer Res 11, 2606–2611 (2005).

63. Grady, W.M. Genomic instability and colon cancer. Cancer Metastasis Rev 23, 11–27 (2004).

64. Ogino, S., Kawasaki, T., Kirkner, G. J., Ohnishi, M. & Fuchs, C. S. 18q loss of heterozygosity in microsatellite stable colorectal cancer is correlated with CpG island methylator phenotype-negative (CIMP-0) and inversely with CIMP-low and CIMP-high. BMC Cancer 7, 72 (2007).

65. Jen, J. et al. Allelic loss of chromosome 18q and prognosis in colorectal cancer. N Engl J Med 331, 213–221 (1994).

66. Jiao, W. et al. A deep learning system accurately classifies primary and metastatic cancers using passenger mutation patterns. Nat Commun 11, 728 (2020).

67. Lorkowski, S. W., Dermawan, J. K. & Rubin, B. P. The practical utility of AI-assisted molecular profiling in the diagnosis and management of cancer of unknown primary: an updated review. Virchows Arch 484, 369–375 (2024).

68. Prendergast, S. C. et al. Sarcoma and the 100,000 Genomes Project: our experience and changes to practice. J Pathol Clin Res 6, 297–307 (2020).

69. Novakovsky, G., Dexter, N., Libbrecht, M. W., Wasserman, W. W. & Mostafavi, S. Obtaining genetics insights from deep learning via explainable artificial intelligence. Nat Rev Genet 24, 125–137 (2023).

70. Turnbull, C. Introducing whole-genome sequencing into routine cancer care: the Genomics England 100 000 Genomes Project. Ann Oncol 29, 784–787 (2018).

71. Sosinsky, A. et al. Insights for precision oncology from the integration of genomic and clinical data of 13,880 tumors from the 100,000 Genomes Cancer Programme. Nat Med 30, 279–289 (2024).

72. Roller, E., Ivakhno, S., Lee, S., Royce, T. & Tanner, S. Canvas: versatile and scalable detection of copy number variants. Bioinformatics 32, 2375–2377 (2016).

73. Gerstung, M. et al. The evolutionary history of 2,658 cancers. Nature 578, 122–128 (2020).

74. Akiba, T., Sano, S., Yanase, T., Ohta, T. & Koyama, M. Optuna: A Next-generation Hyperparameter Optimization Framework. Preprint at 10.48550/arXiv.1907.10902 (2019).

75. Paszke, A. et al. PyTorch: An Imperative Style, High-Performance Deep Learning Library. Preprint at 10.48550/arXiv.1912.01703 (2019).

76. Jha, A., K Aicher, J., R Gazzara, M., Singh, D. & Barash, Y. Enhanced Integrated Gradients: improving interpretability of deep learning models using splicing codes as a case study. Genome Biol 21, 149 (2020).

77. Kassambara, A., Kosinski, M., Biecek, P. & Fabian, S. survminer: Drawing Survival Curves using ‘ggplot2’. (2021).

