## Supplementary Information for "DeepCNA: an explainable deep learning method for cancer diagnosis and cancer-specific patterns of copy number aberrations"

### Contents

- Figures 1-13: Classification probability boxplots
- Figures 14-26: Circular Manhattan plots
- Figures 27-39: Bubble plots for chromosome arm attribute
- Figures 40-65: Heatmaps of high attribute CNAs

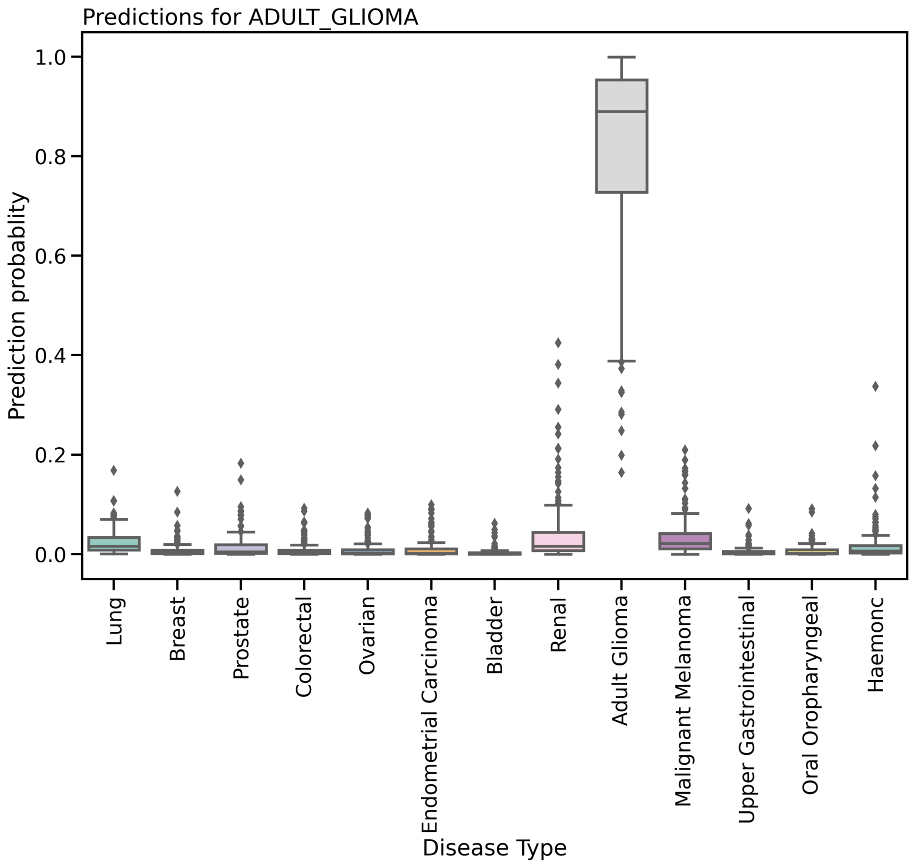


Supplementary Figure 1: Classification probability boxplots for the Adult Glioma cohort across the 13 cancer primary sites for the WGS classification task.

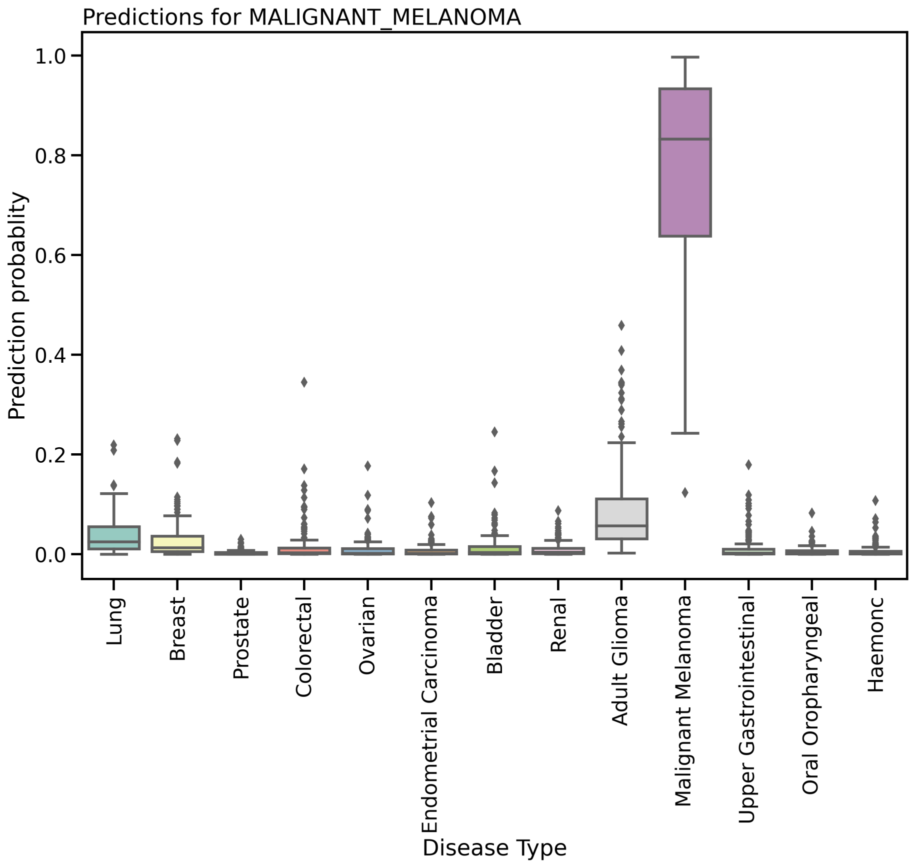


Supplementary Figure 2: Classification probability boxplots for the Malignant Melanoma cohort across the 13 cancer primary sites for the WGS classification task.

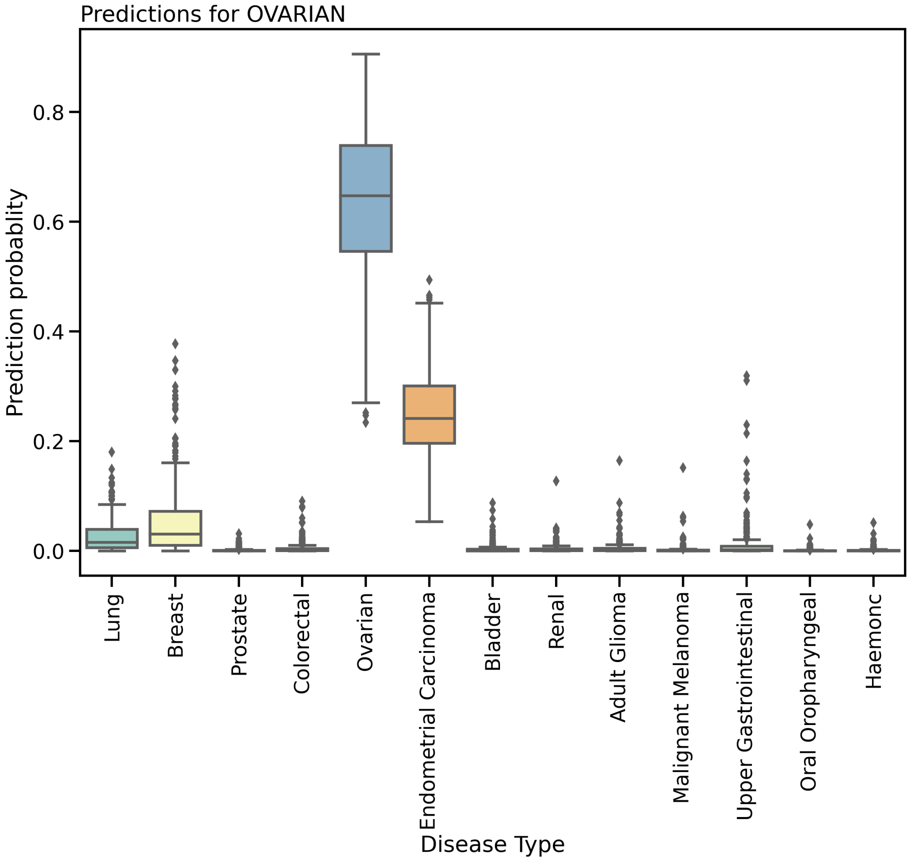


Supplementary Figure 3: Classification probability boxplots for the Ovarian cohort across the 13 cancer primary sites for the WGS classification task.

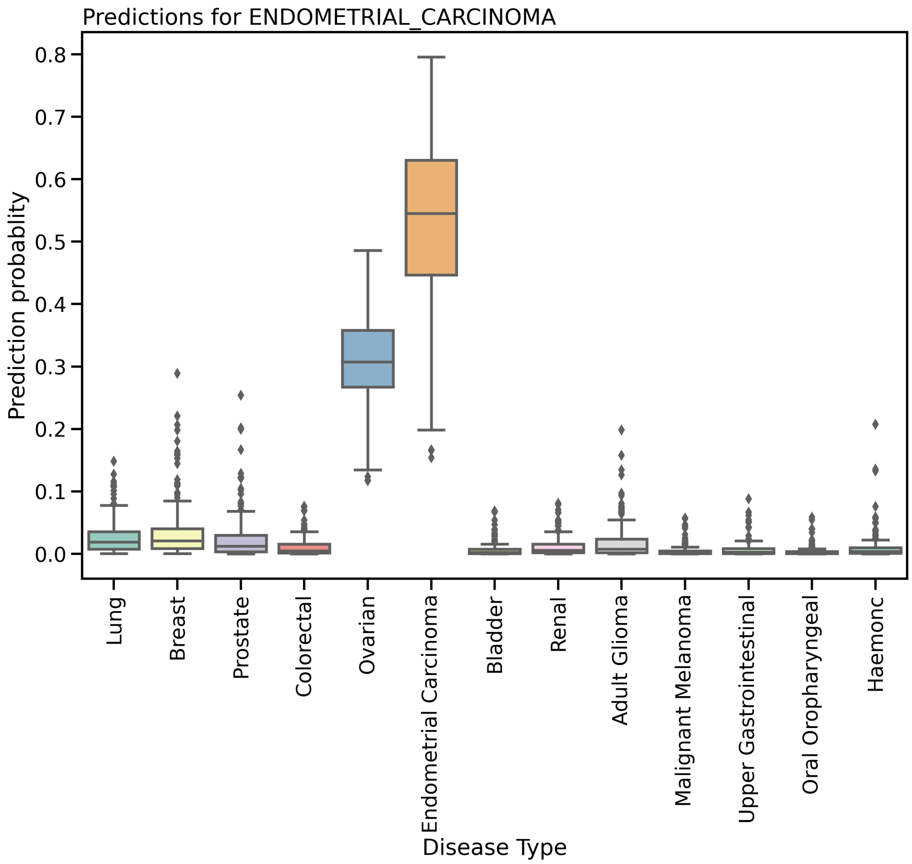


Supplementary Figure 4: Classification probability boxplots for the Endometrial Carcinoma cohort across the 13 cancer primary sites for the WGS classification task.

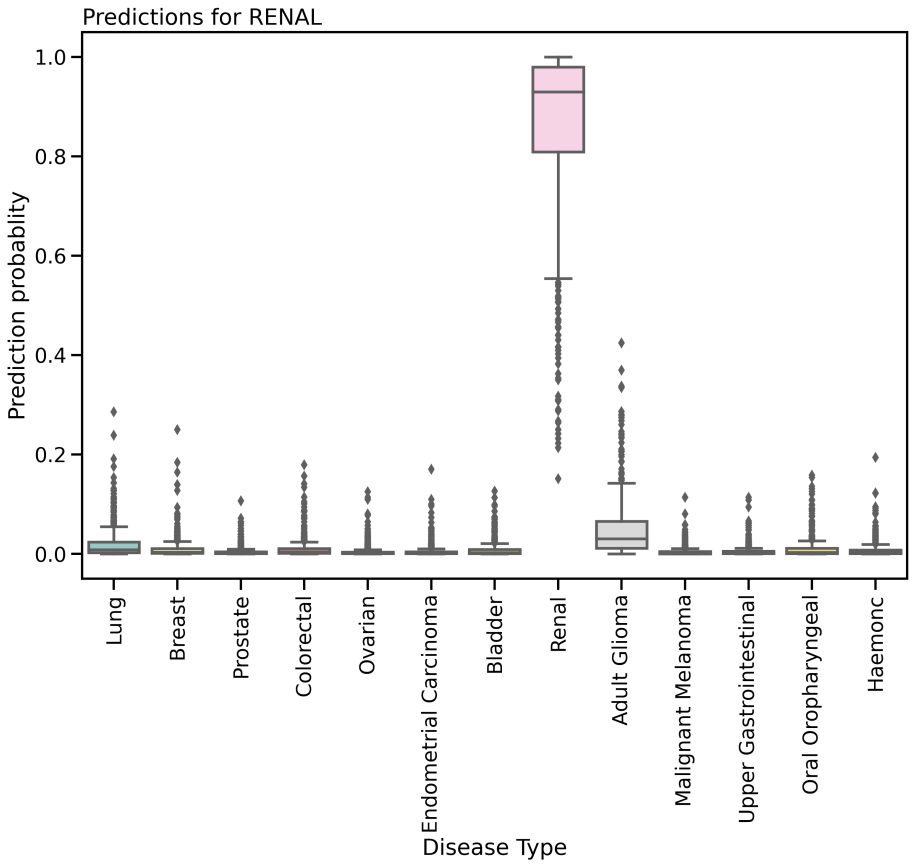


Supplementary Figure 5: Classification probability boxplots for the Renal cohort across the 13 cancer primary sites for the WGS classification task.

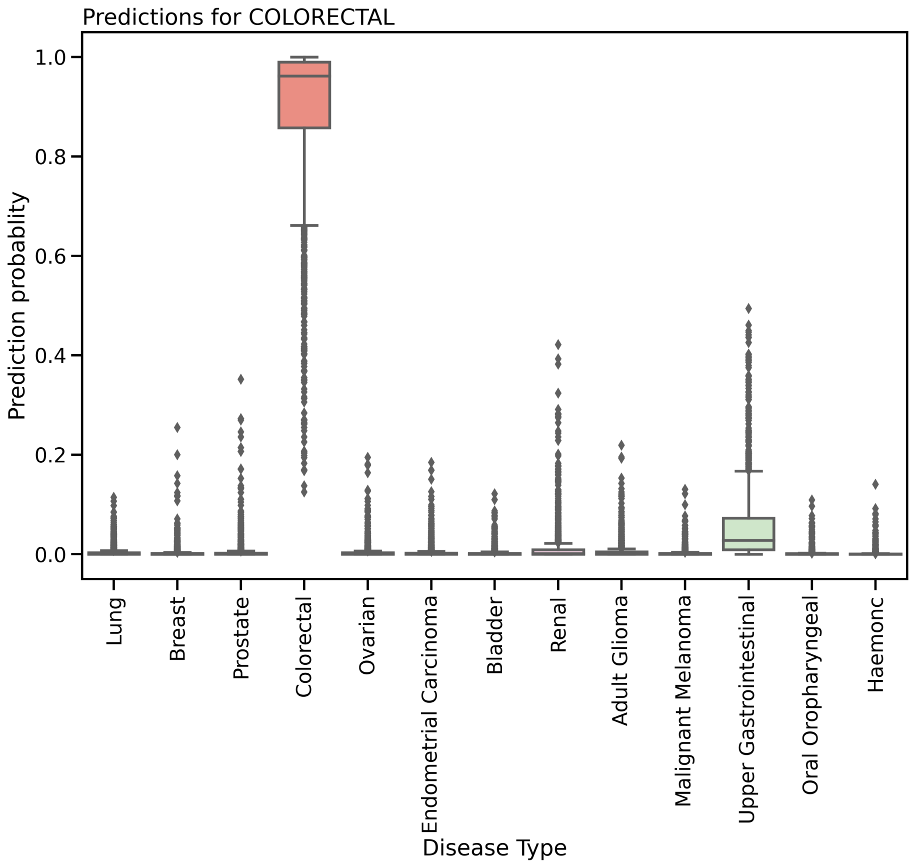


Supplementary Figure 6: Classification probability boxplots for the Colorectal cohort across the 13 cancer primary sites for the WGS classification task.

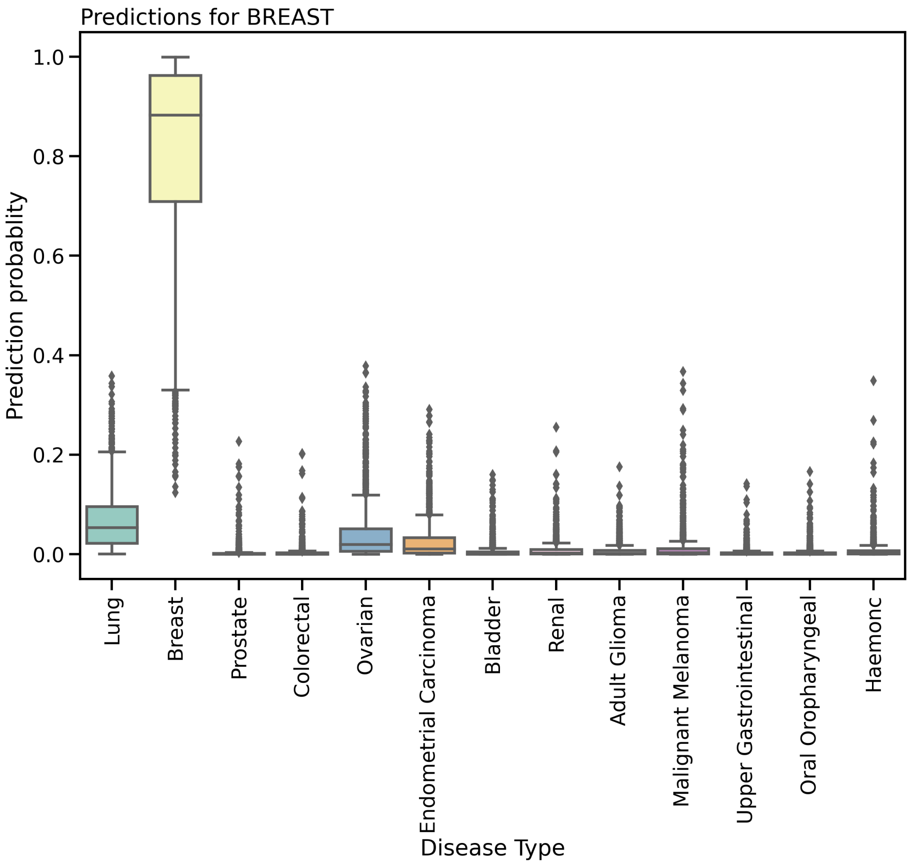


Supplementary Figure 7: Classification probability boxplots for the Breast cohort across the 13 cancer primary sites for the WGS classification task.

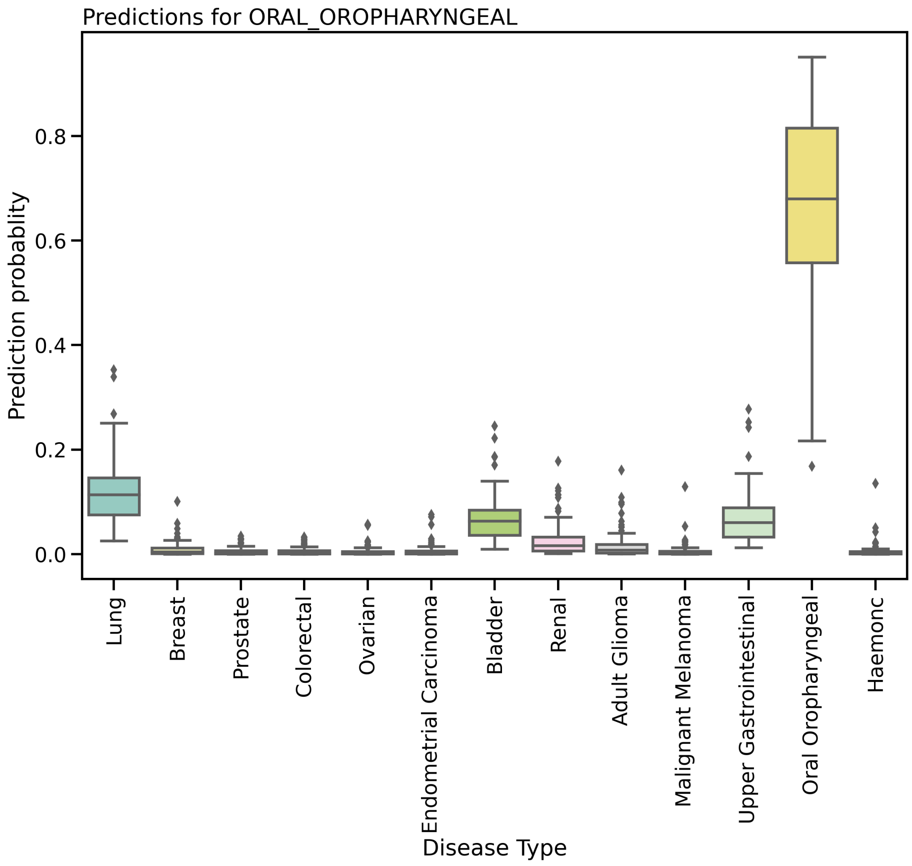


Supplementary Figure 8: Classification probability boxplots for the Oral Oropharyngeal cohort across the 13 cancer primary sites for the WGS classification task.

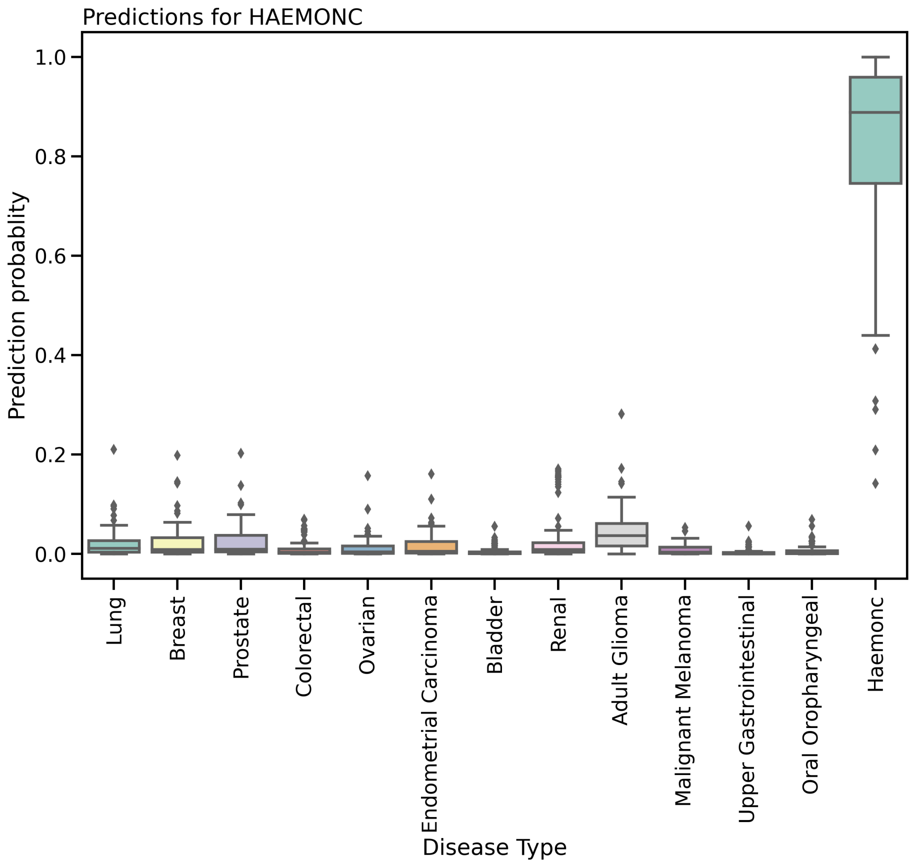


Supplementary Figure 9: Classification probability boxplots for the Haemonc cohort across the 13 cancer primary sites for the WGS classification task.

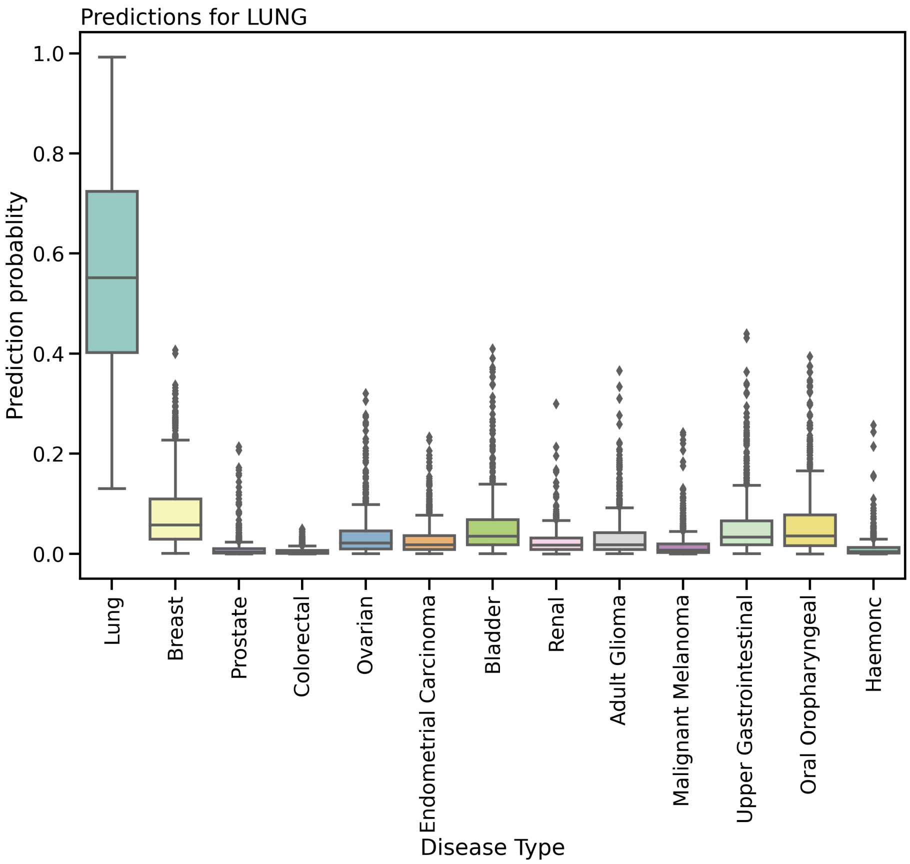


Supplementary Figure 10: Classification probability boxplots for the Lung cohort across the 13 cancer primary sites for the WGS classification task.

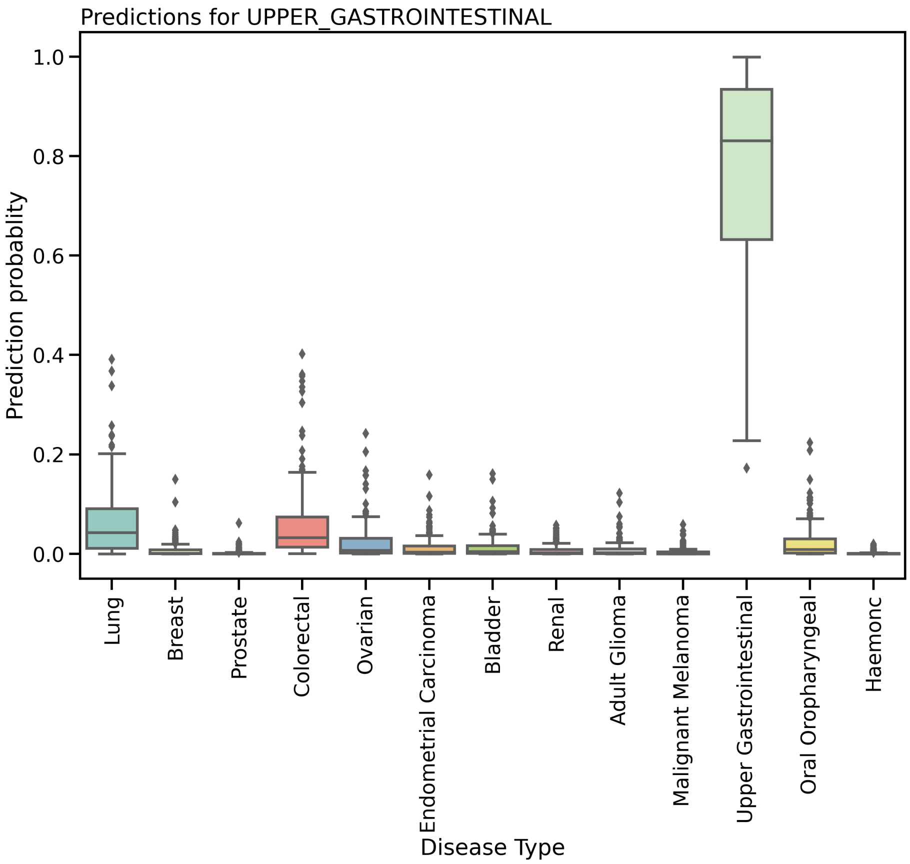


Supplementary Figure 11: Classification probability boxplots for the Upper Gastrointestinal cohort across the 13 cancer primary sites for the WGS classification task.

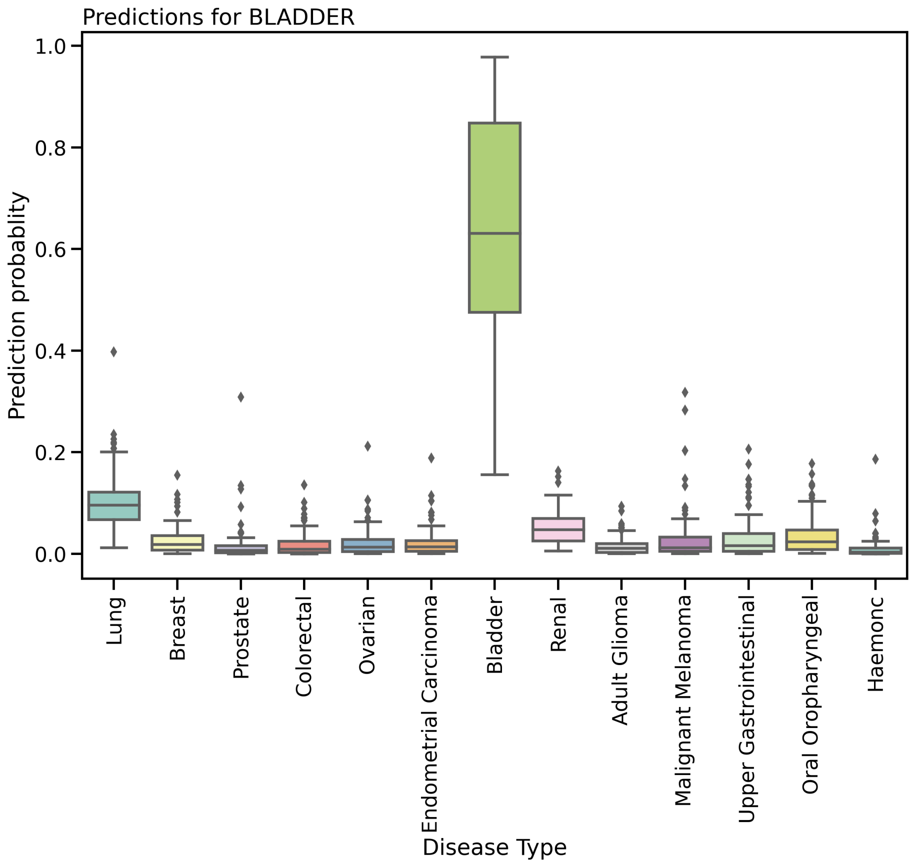


Supplementary Figure 12: Classification probability boxplots for the Bladder cohort across the 13 cancer primary sites for the WGS classification task.

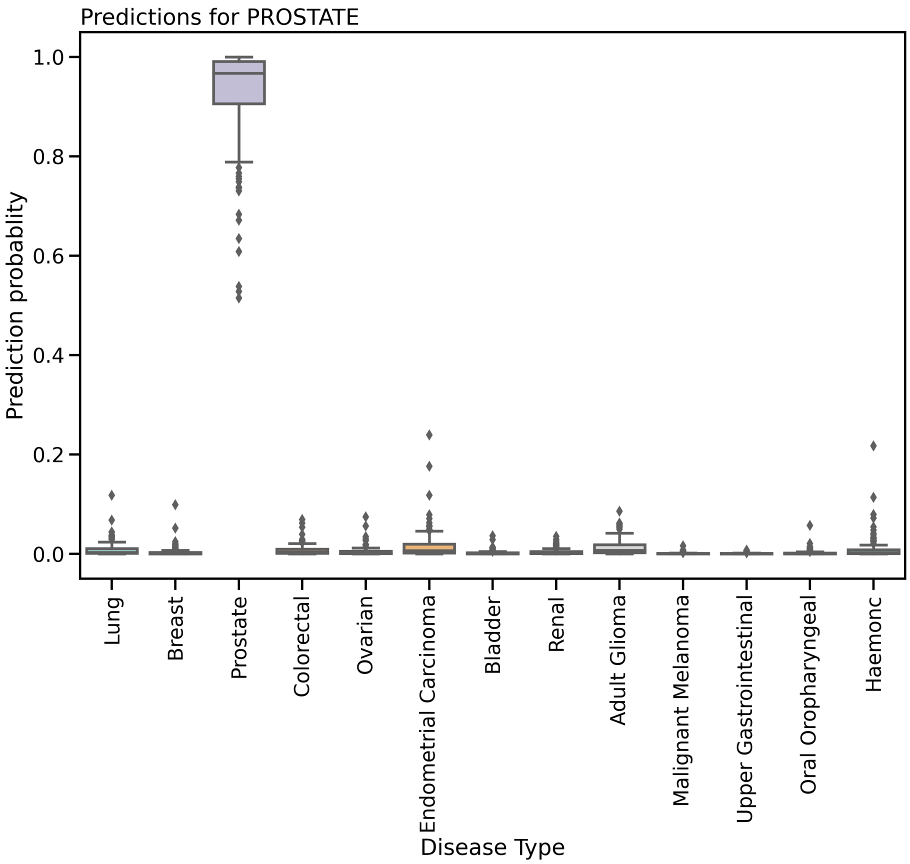


Supplementary Figure 13: Classification probability boxplots for the Prostate cohort across the 13 cancer primary sites for the WGS classification task.

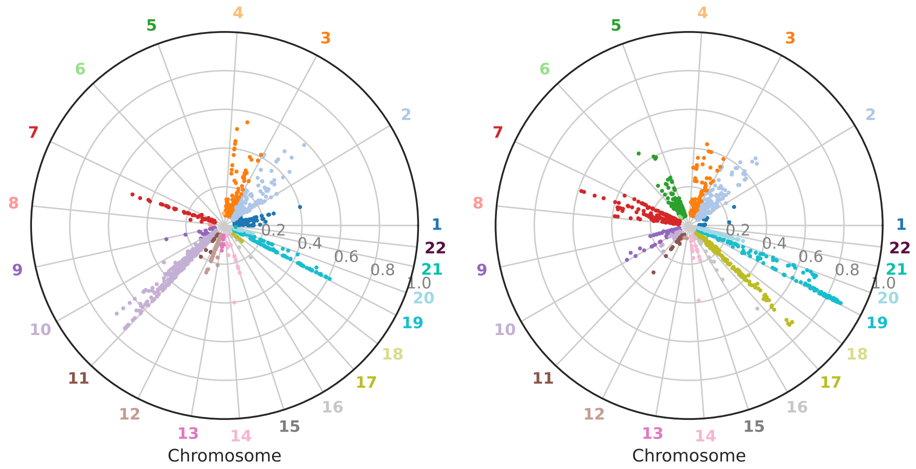


Supplementary Figure 14: Circular Manhattan Plot for Endometrial Carcinoma cohort for total allele copy number (left) and minor allele copy number (right)

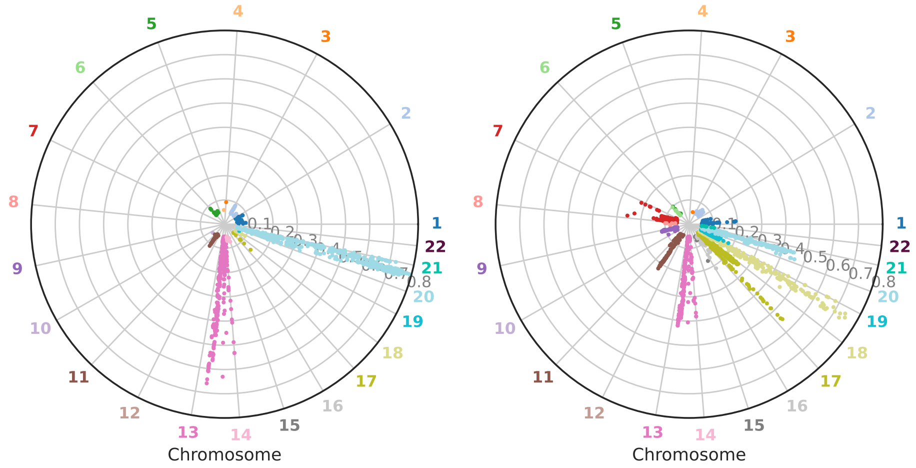


Supplementary Figure 15: Circular Manhattan Plot for Colorectal cohort for total allele copy number (left) and minor allele copy number (right)

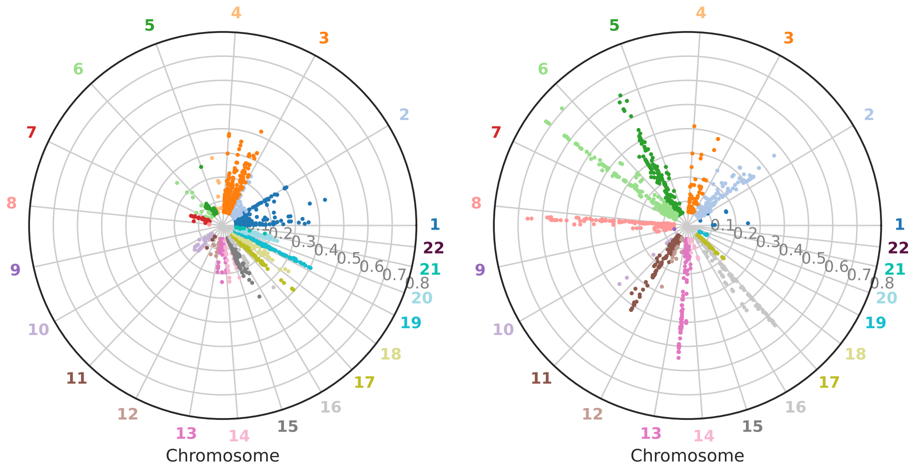


Supplementary Figure 16: Circular Manhattan Plot for Prostate cohort for total allele copy number (left) and minor allele copy number (right)

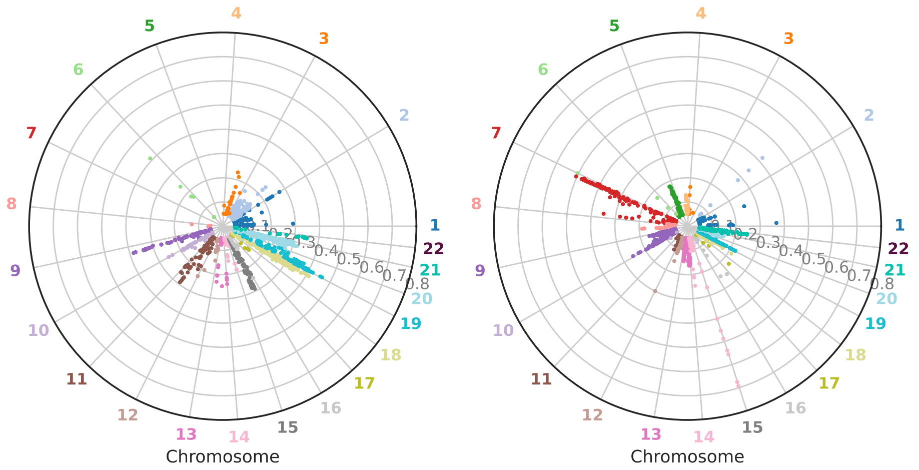


Supplementary Figure 17: Circular Manhattan Plot for Haemonc cohort for total allele copy number (left) and minor allele copy number (right)

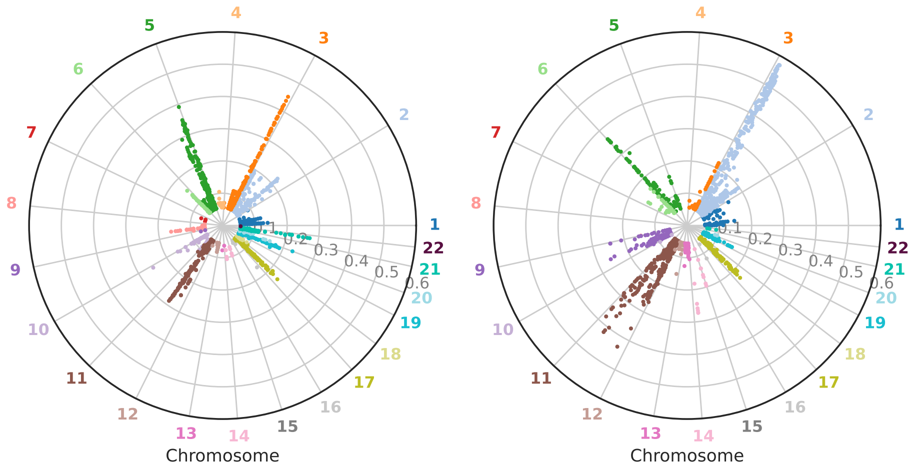


Supplementary Figure 18: Circular Manhattan Plot for Bladder cohort for total allele copy number (left) and minor allele copy number (right)

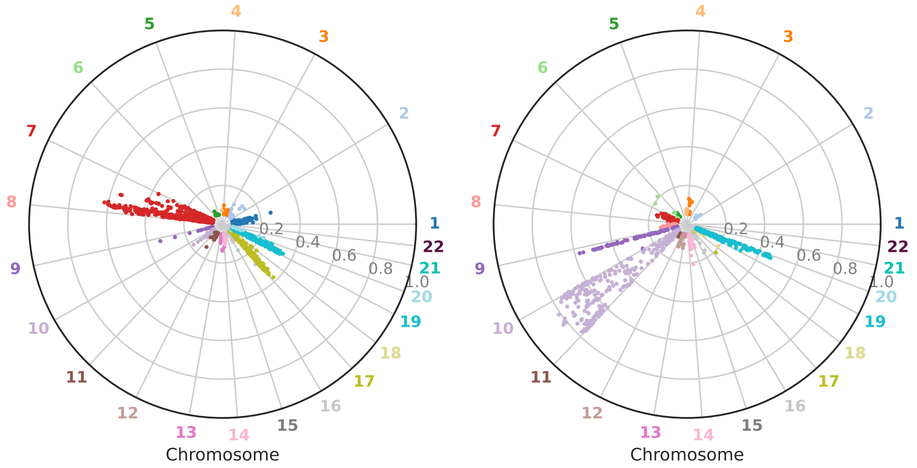


Supplementary Figure 19: Circular Manhattan Plot for Adult Glioma cohort for total allele copy number (left) and minor allele copy number (right)

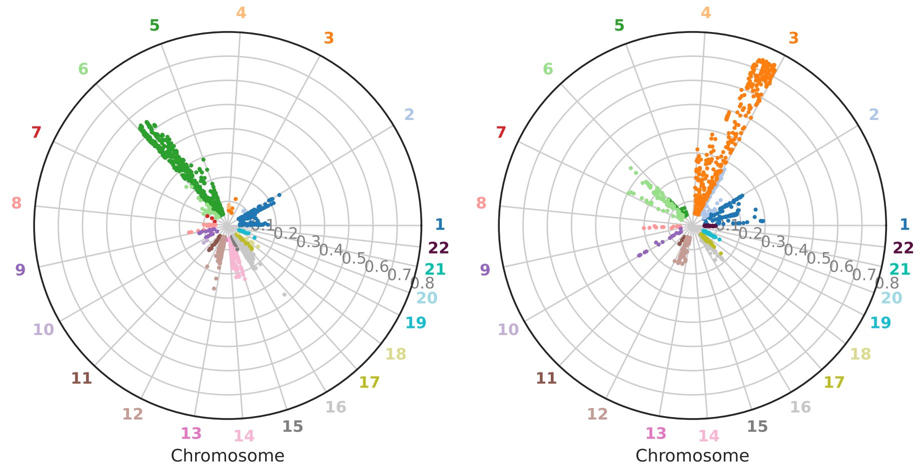


Supplementary Figure 20: Circular Manhattan Plot for Renal cohort for total allele copy number (left) and minor allele copy number (right)

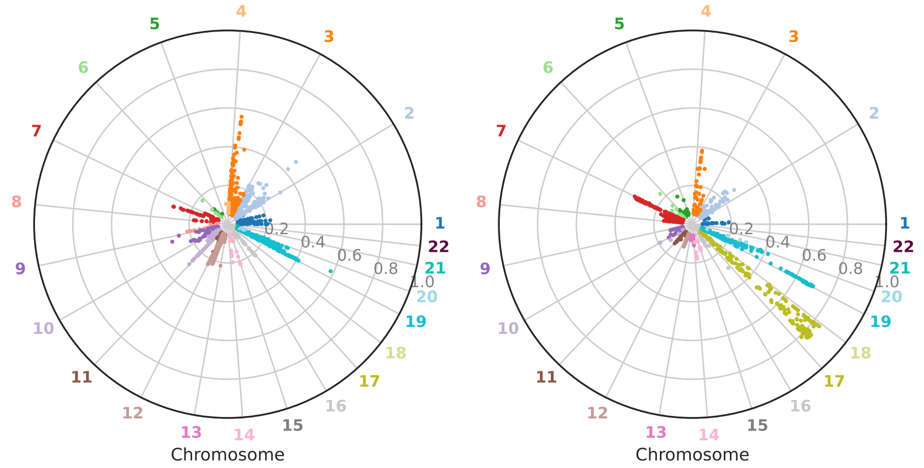


Supplementary Figure 21: Circular Manhattan Plot for Ovarian cohort for total allele copy number (left) and minor allele copy number (right)

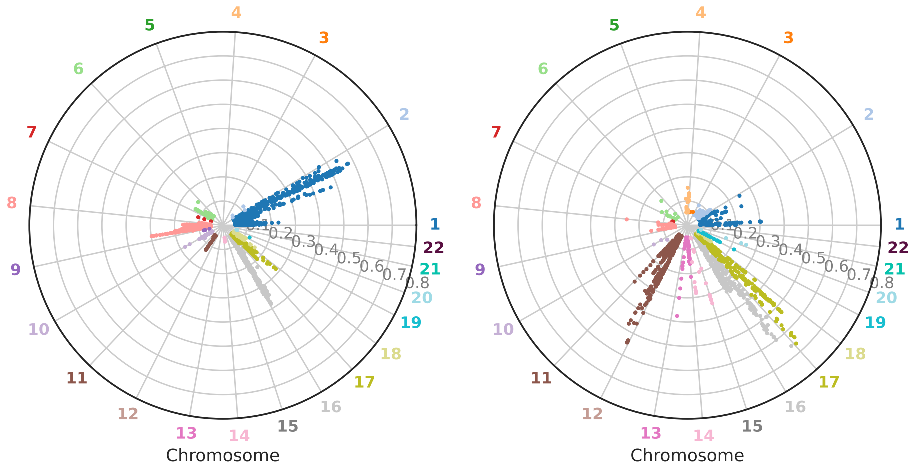


Supplementary Figure 22: Circular Manhattan Plot for Breast cohort for total allele copy number (left) and minor allele copy number (right)

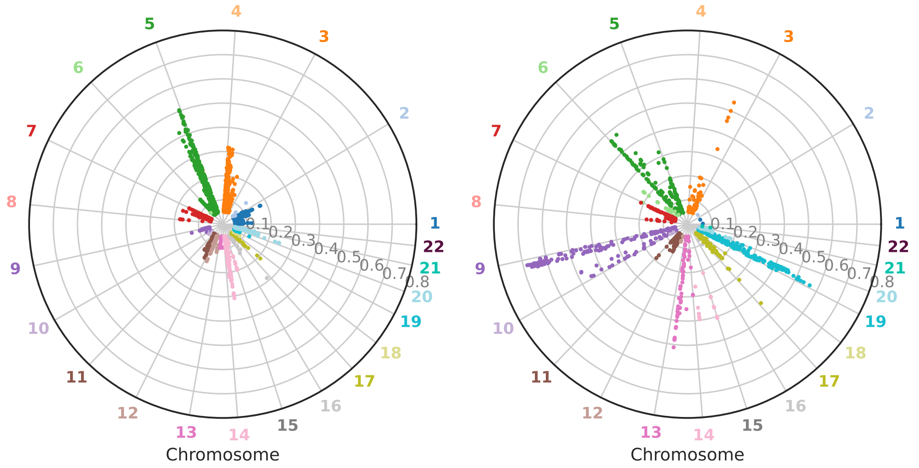


Supplementary Figure 23: Circular Manhattan Plot for Lung cohort for total allele copy number (left) and minor allele copy number (right)

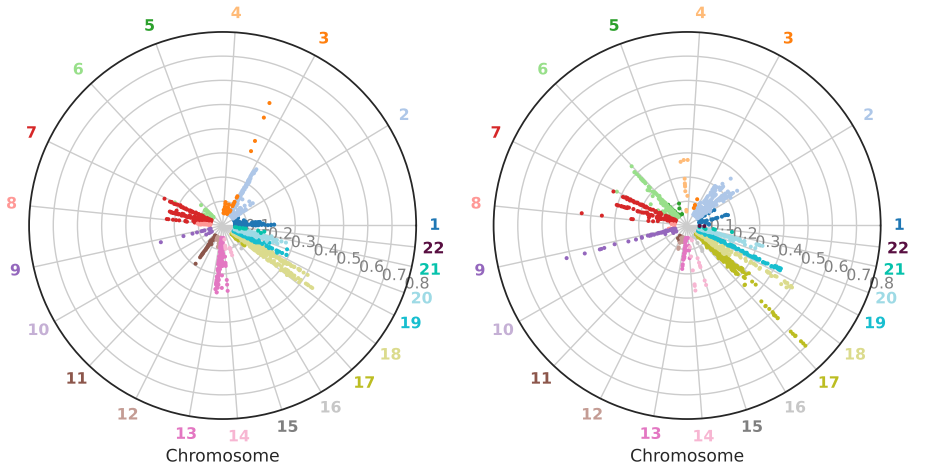


Supplementary Figure 24: Circular Manhattan Plot for Upper Gastrointestinal cohort for total allele copy number (left) and minor allele copy number (right)

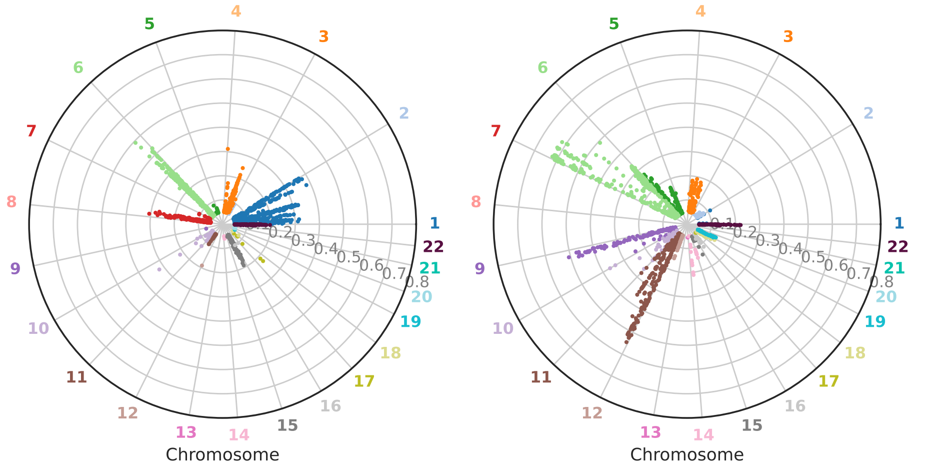


Supplementary Figure 25: Circular Manhattan Plot for Malignant Melanoma cohort for total allele copy number (left) and minor allele copy number (right)

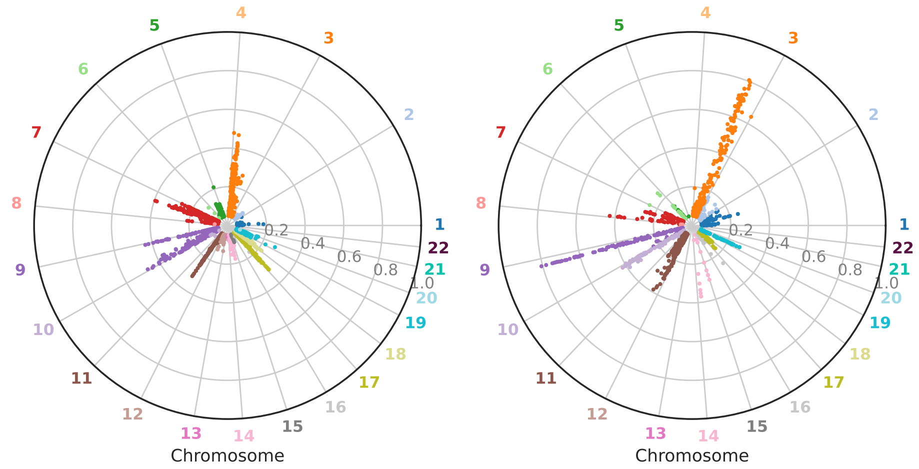


Supplementary Figure 26: Circular Manhattan Plot for Oral Oropharyngeal cohort for total allele copy number (left) and minor allele copy number (right)

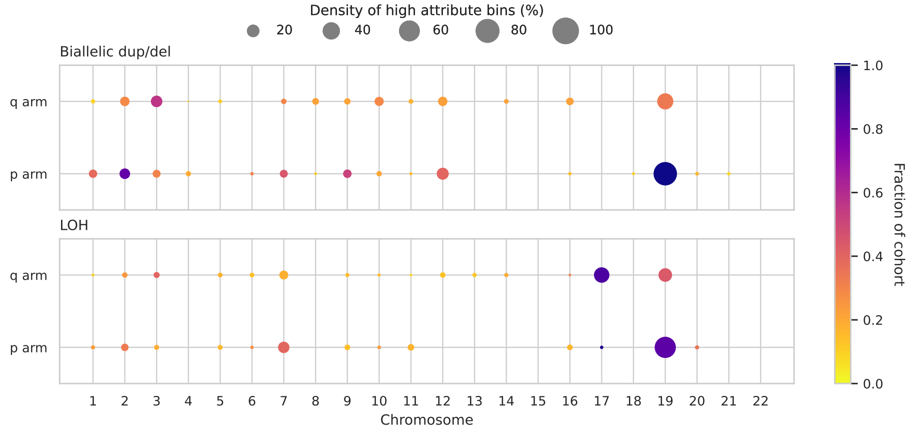


Supplementary Figure 27: Bubble plots for Ovarian cohort for both total and minor allele copy numbers, displaying high-attribute loci across chromosome arms. Bubble size represents the density of significant bins, with colour indicating the fraction of the cohort sharing these loci.

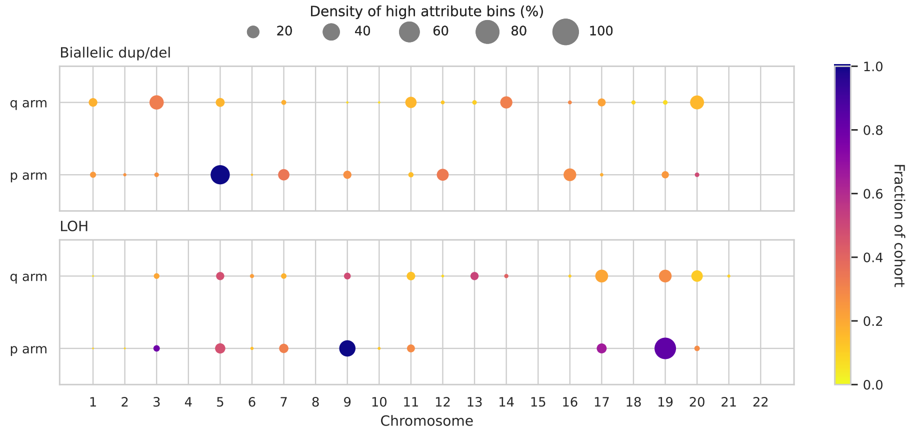


Supplementary Figure 28: Bubble plots for Lung cohort for both total and minor allele copy numbers, displaying high-attribute loci across chromosome arms. Bubble size represents the density of significant bins, with colour indicating the fraction of the cohort sharing these loci.

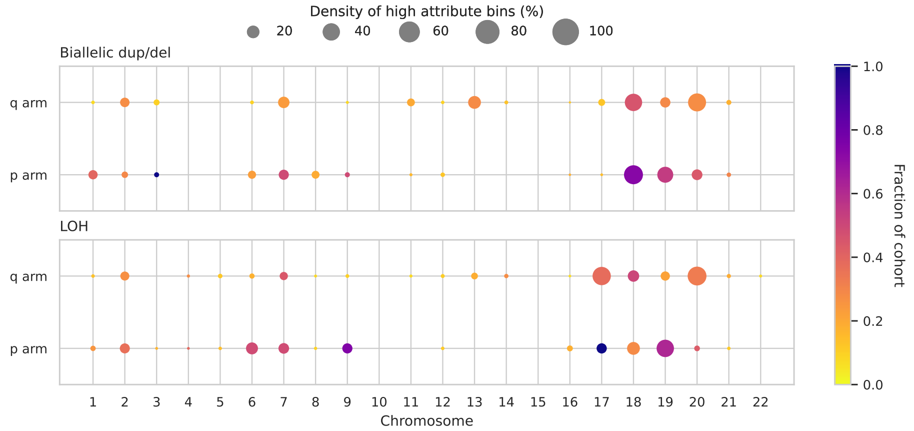


Supplementary Figure 29: Bubble plots for Upper Gastrointestinal cohort for both total and minor allele copy numbers, displaying high-attribute loci across chromosome arms. Bubble size represents the density of significant bins, with colour indicating the fraction of the cohort sharing these loci.

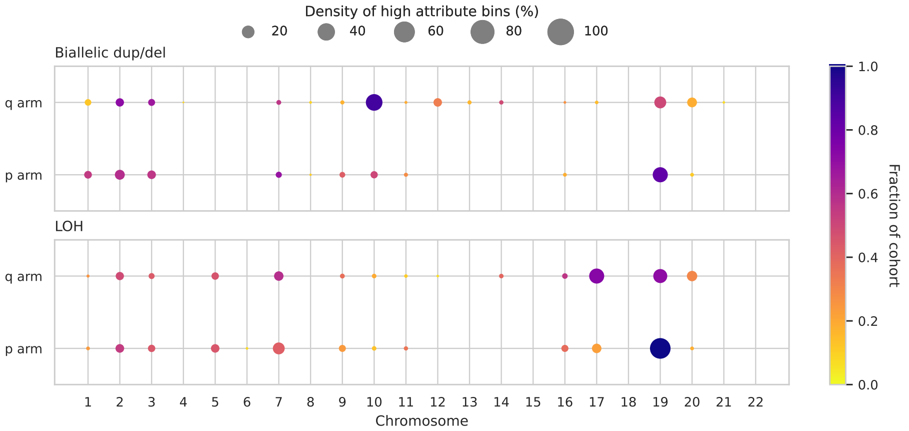


Supplementary Figure 30: Bubble plots for Endometrial Carcinoma cohort for both total and minor allele copy numbers, displaying high-attribute loci across chromosome arms. Bubble size represents the density of significant bins, with colour indicating the fraction of the cohort sharing these loci.

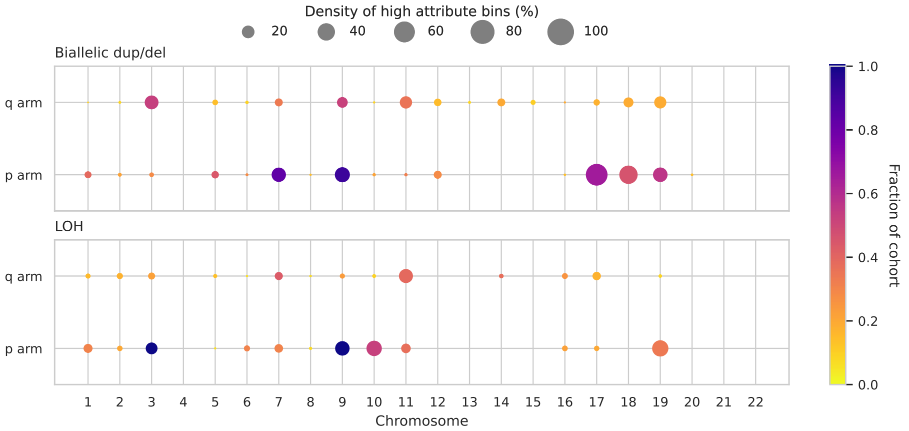


Supplementary Figure 31: Bubble plots for Oral Oropharyngeal cohort for both total and minor allele copy numbers, displaying high-attribute loci across chromosome arms. Bubble size represents the density of significant bins, with colour indicating the fraction of the cohort sharing these loci.

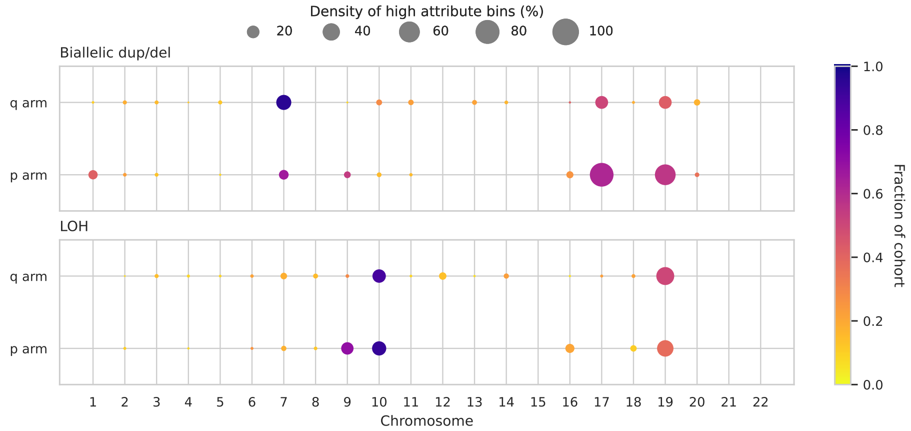


Supplementary Figure 32: Bubble plots for Adult Glioma cohort for both total and minor allele copy numbers, displaying high-attribute loci across chromosome arms. Bubble size represents the density of significant bins, with colour indicating the fraction of the cohort sharing these loci.

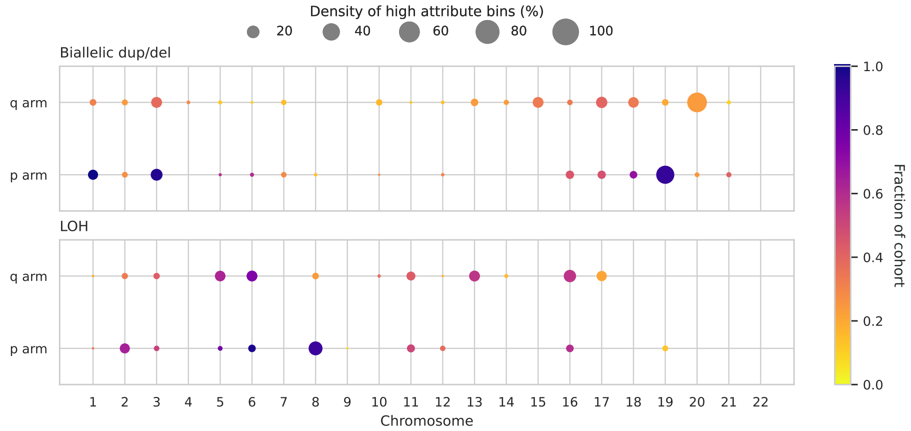


Supplementary Figure 33: Bubble plots for Prostate cohort for both total and minor allele copy numbers, displaying high-attribute loci across chromosome arms. Bubble size represents the density of significant bins, with colour indicating the fraction of the cohort sharing these loci.

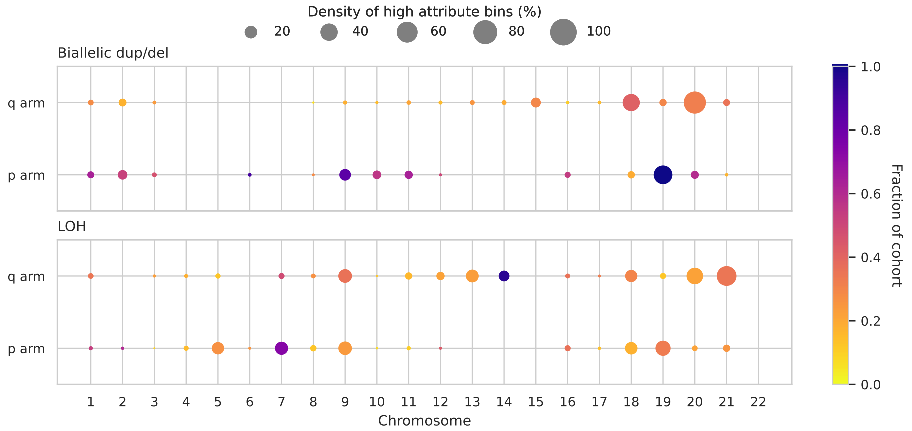


Supplementary Figure 34: Bubble plots for Haemonc cohort for both total and minor allele copy numbers, displaying high-attribute loci across chromosome arms. Bubble size represents the density of significant bins, with colour indicating the fraction of the cohort sharing these loci.

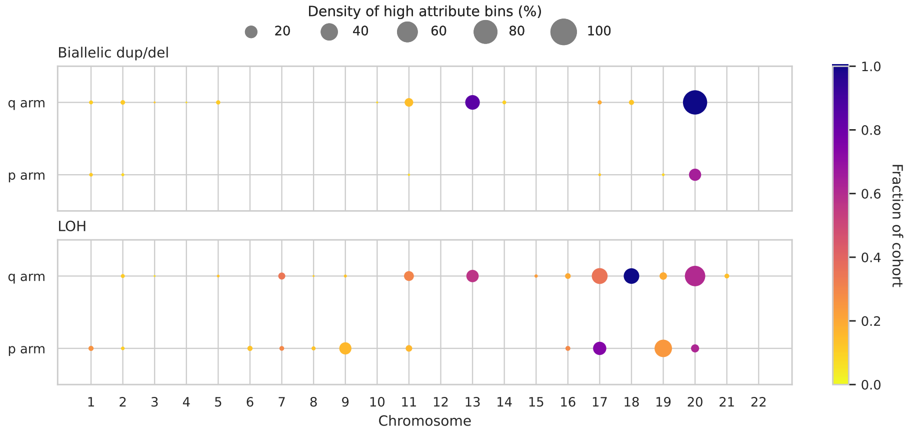


Supplementary Figure 35: Bubble plots for Colorectal cohort for both total and minor allele copy numbers, displaying high-attribute loci across chromosome arms. Bubble size represents the density of significant bins, with colour indicating the fraction of the cohort sharing these loci.

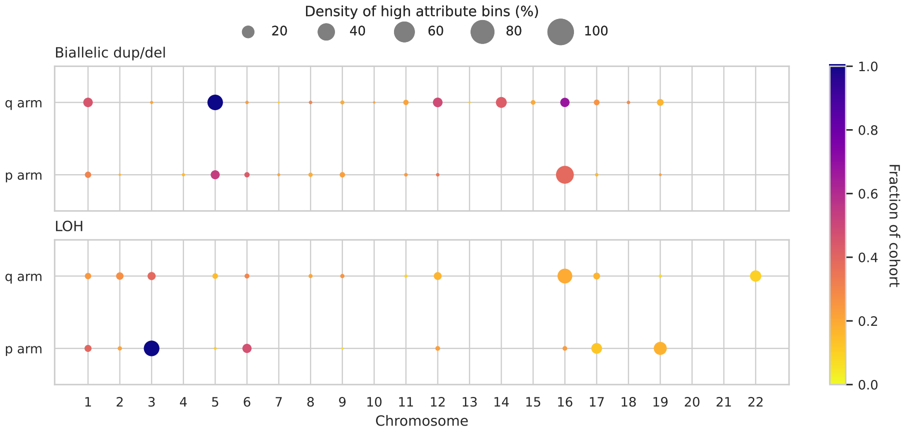


Supplementary Figure 36: Bubble plots for Renal cohort for both total and minor allele copy numbers, displaying high-attribute loci across chromosome arms. Bubble size represents the density of significant bins, with colour indicating the fraction of the cohort sharing these loci.

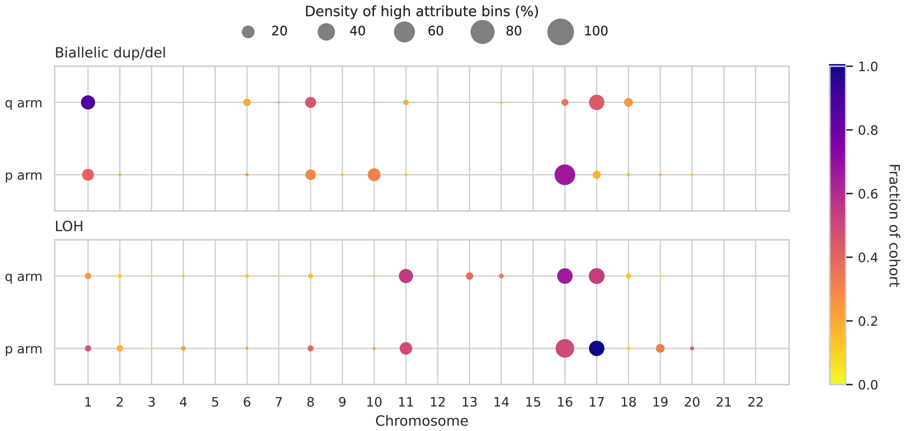


Supplementary Figure 37: Bubble plots for Breast cohort for both total and minor allele copy numbers, displaying high-attribute loci across chromosome arms. Bubble size represents the density of significant bins, with colour indicating the fraction of the cohort sharing these loci.

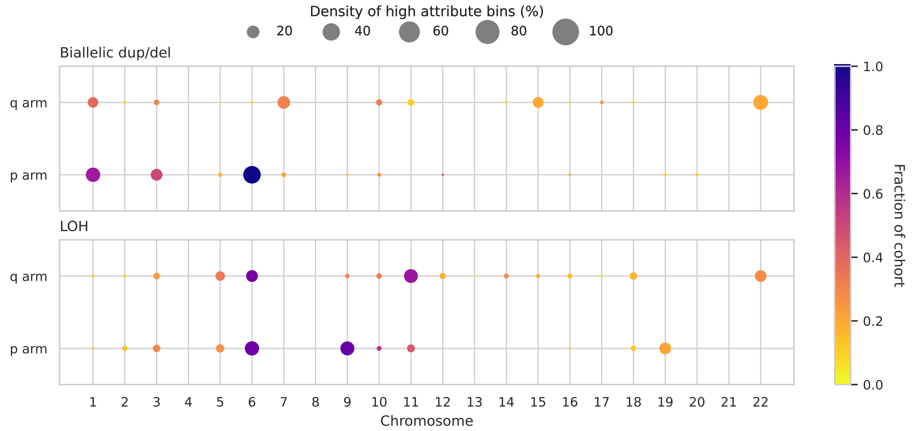


Supplementary Figure 38: Bubble plots for Malignant Melanoma cohort for both total and minor allele copy numbers, displaying high-attribute loci across chromosome arms. Bubble size represents the density of significant bins, with colour indicating the fraction of the cohort sharing these loci.

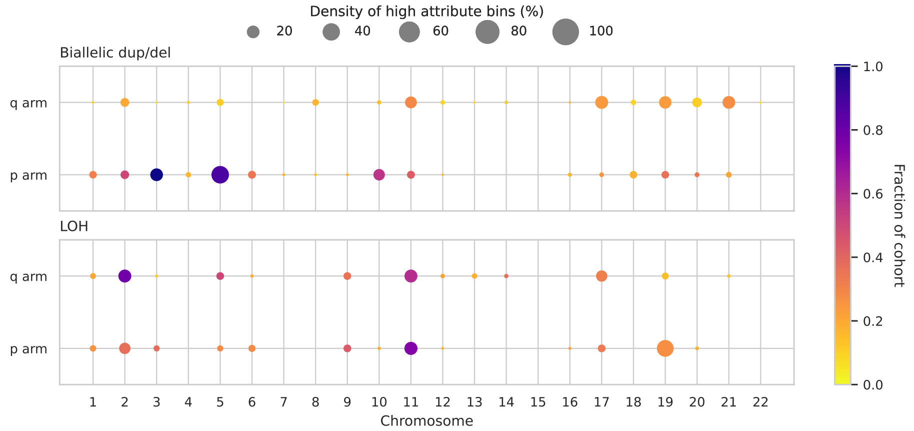


Supplementary Figure 39: Bubble plots for Bladder cohort for both total and minor allele copy numbers, displaying high-attribute loci across chromosome arms. Bubble size represents the density of significant bins, with colour indicating the fraction of the cohort sharing these loci.

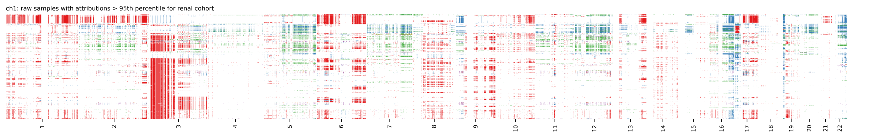


Supplementary Figure 40: Heatmap of CNA for samples in Renal cohort for minor allele copy number. It highlights only high attribute loci as determined by the model, coloured according to CNA values. Loci not selected by the model are omitted, appearing as white spaces, to focus on regions of highest model attribution.

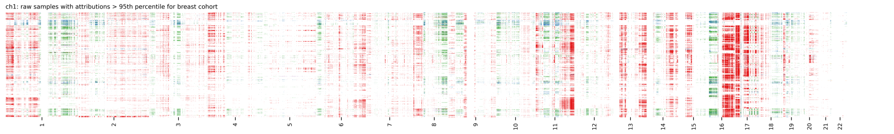


Supplementary Figure 41: Heatmap of CNA for samples in Breast cohort for minor allele copy number. It highlights only high attribute loci as determined by the model, coloured according to CNA values. Loci not selected by the model are omitted, appearing as white spaces, to focus on regions of highest model attribution.

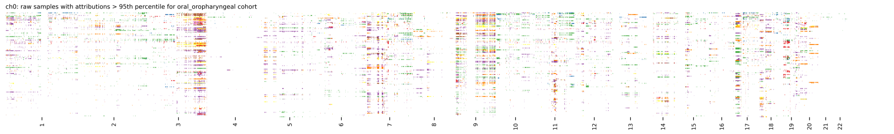


Supplementary Figure 42: Heatmap of CNA for samples in Oral Oropharyngeal cohort for total allele copy number. It highlights only high attribute loci as determined by the model, coloured according to CNA values. Loci not selected by the model are omitted, appearing as white spaces, to focus on regions of highest model attribution.

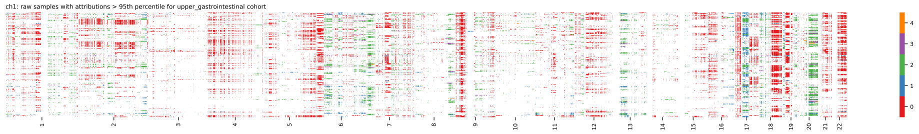


Supplementary Figure 43: Heatmap of CNA for samples in Upper Gastrointestinal cohort for minor allele copy number. It highlights only high attribute loci as determined by the model, coloured according to CNA values. Loci not selected by the model are omitted, appearing as white spaces, to focus on regions of highest model attribution.

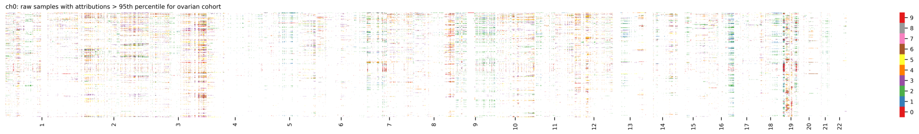


Supplementary Figure 44: Heatmap of CNA for samples in Ovarian cohort for total allele copy number. It highlights only high attribute loci as determined by the model, coloured according to CNA values. Loci not selected by the model are omitted, appearing as white spaces, to focus on regions of highest model attribution.

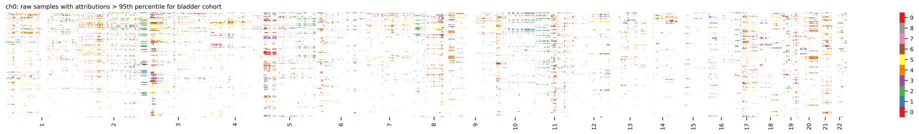


Supplementary Figure 45: Heatmap of CNA for samples in Bladder cohort for total allele copy number. It highlights only high attribute loci as determined by the model, coloured according to CNA values. Loci not selected by the model are omitted, appearing as white spaces, to focus on regions of highest model attribution.

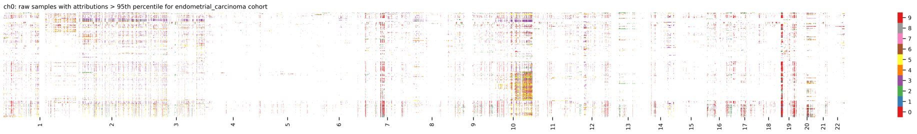


Supplementary Figure 46: Heatmap of CNA for samples in Endometrial Carcinoma cohort for total allele copy number. It highlights only high attribute loci as determined by the model, coloured according to CNA values. Loci not selected by the model are omitted, appearing as white spaces, to focus on regions of highest model attribution.

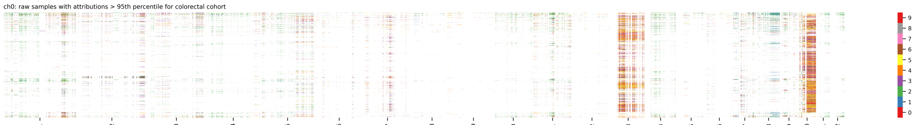


Supplementary Figure 47: Heatmap of CNA for samples in Colorectal cohort for total allele copy number. It highlights only high attribute loci as determined by the model, coloured according to CNA values. Loci not selected by the model are omitted, appearing as white spaces, to focus on regions of highest model attribution.

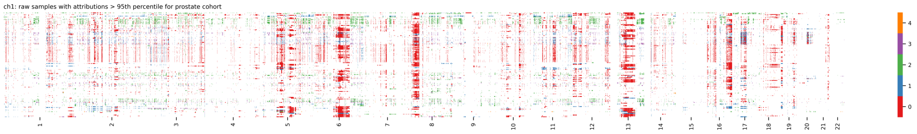


Supplementary Figure 48: Heatmap of CNA for samples in Prostate cohort for minor allele copy number. It highlights only high attribute loci as determined by the model, coloured according to CNA values. Loci not selected by the model are omitted, appearing as white spaces, to focus on regions of highest model attribution.

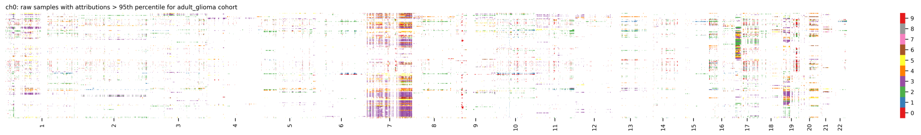


Supplementary Figure 49: Heatmap of CNA for samples in Adult Glioma cohort for total allele copy number. It highlights only high attribute loci as determined by the model, coloured according to CNA values. Loci not selected by the model are omitted, appearing as white spaces, to focus on regions of highest model attribution.

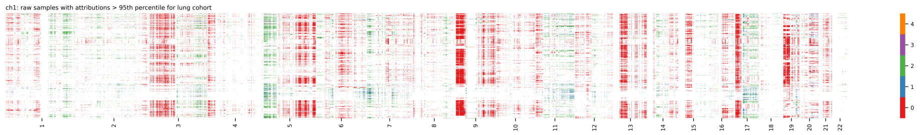


Supplementary Figure 50: Heatmap of CNA for samples in Lung cohort for minor allele copy number. It highlights only high attribute loci as determined by the model, coloured according to CNA values. Loci not selected by the model are omitted, appearing as white spaces, to focus on regions of highest model attribution.

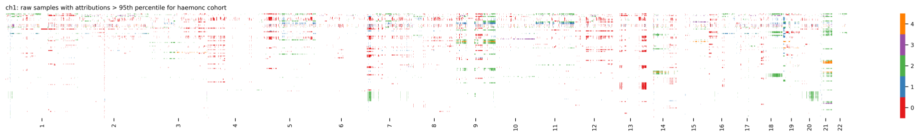


Supplementary Figure 51: Heatmap of CNA for samples in Haemonc cohort for minor allele copy number. It highlights only high attribute loci as determined by the model, coloured according to CNA values. Loci not selected by the model are omitted, appearing as white spaces, to focus on regions of highest model attribution.

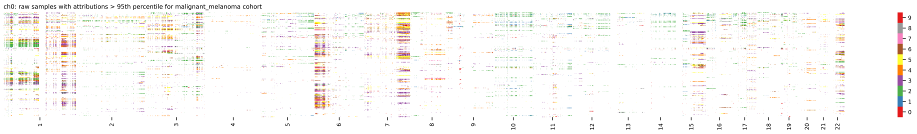


Supplementary Figure 52: Heatmap of CNA for samples in Malignant Melanoma cohort for total allele copy number. It highlights only high attribute loci as determined by the model, coloured according to CNA values. Loci not selected by the model are omitted, appearing as white spaces, to focus on regions of highest model attribution.

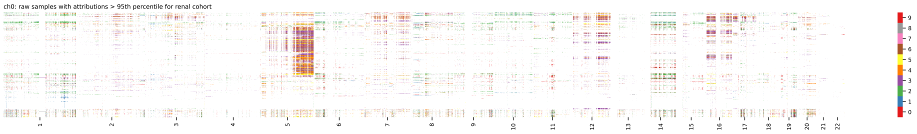


Supplementary Figure 53: Heatmap of CNA for samples in Renal cohort for total allele copy number. It highlights only high attribute loci as determined by the model, coloured according to CNA values. Loci not selected by the model are omitted, appearing as white spaces, to focus on regions of highest model attribution.

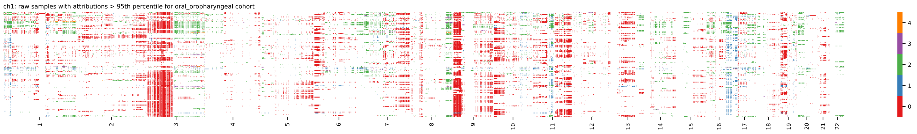


Supplementary Figure 54: Heatmap of CNA for samples in Oral Oropharyngeal cohort for minor allele copy number. It highlights only high attribute loci as determined by the model, coloured according to CNA values. Loci not selected by the model are omitted, appearing as white spaces, to focus on regions of highest model attribution.

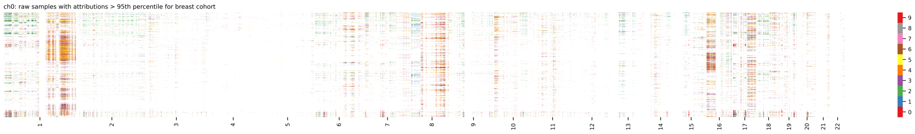


Supplementary Figure 55: Heatmap of CNA for samples in Breast cohort for total allele copy number. It highlights only high attribute loci as determined by the model, coloured according to CNA values. Loci not selected by the model are omitted, appearing as white spaces, to focus on regions of highest model attribution.

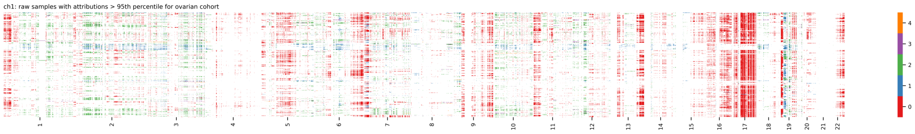


Supplementary Figure 56: Heatmap of CNA for samples in Ovarian cohort for minor allele copy number. It highlights only high attribute loci as determined by the model, coloured according to CNA values. Loci not selected by the model are omitted, appearing as white spaces, to focus on regions of highest model attribution.

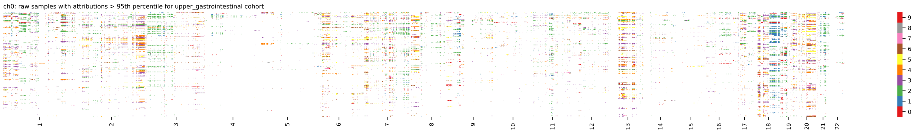


Supplementary Figure 57: Heatmap of CNA for samples in Upper Gastrointestinal cohort for total allele copy number. It highlights only high attribute loci as determined by the model, coloured according to CNA values. Loci not selected by the model are omitted, appearing as white spaces, to focus on regions of highest model attribution.

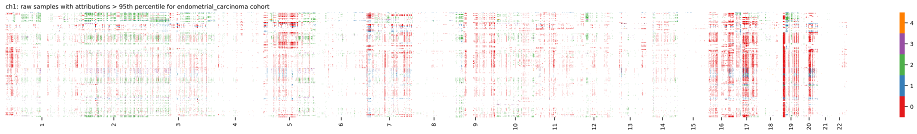


Supplementary Figure 58: Heatmap of CNA for samples in Endometrial Carcinoma cohort for minor allele copy number. It highlights only high attribute loci as determined by the model, coloured according to CNA values. Loci not selected by the model are omitted, appearing as white spaces, to focus on regions of highest model attribution.

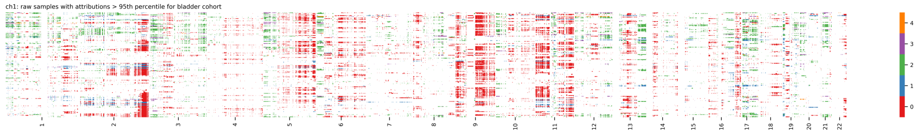


Supplementary Figure 59: Heatmap of CNA for samples in Bladder cohort for minor allele copy number. It highlights only high attribute loci as determined by the model, coloured according to CNA values. Loci not selected by the model are omitted, appearing as white spaces, to focus on regions of highest model attribution.

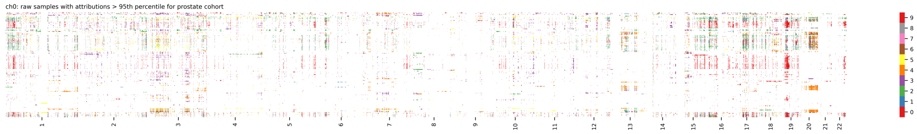


Supplementary Figure 60: Heatmap of CNA for samples in Prostate cohort for total allele copy number. It highlights only high attribute loci as determined by the model, coloured according to CNA values. Loci not selected by the model are omitted, appearing as white spaces, to focus on regions of highest model attribution.

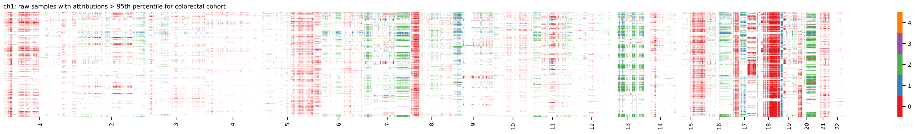


Supplementary Figure 61: Heatmap of CNA for samples in Colorectal cohort for minor allele copy number. It highlights only high attribute loci as determined by the model, coloured according to CNA values. Loci not selected by the model are omitted, appearing as white spaces, to focus on regions of highest model attribution.

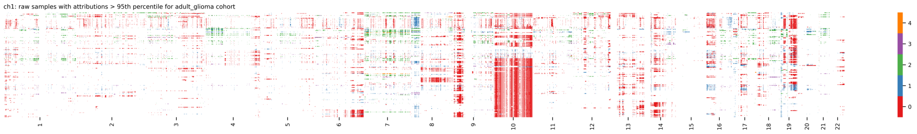


Supplementary Figure 62: Heatmap of CNA for samples in Adult Glioma cohort for minor allele copy number. It highlights only high attribute loci as determined by the model, coloured according to CNA values. Loci not selected by the model are omitted, appearing as white spaces, to focus on regions of highest model attribution.

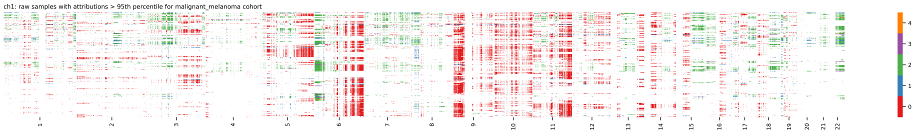


Supplementary Figure 63: Heatmap of CNA for samples in Malignant Melanoma cohort for minor allele copy number. It highlights only high attribute loci as determined by the model, coloured according to CNA values. Loci not selected by the model are omitted, appearing as white spaces, to focus on regions of highest model attribution.

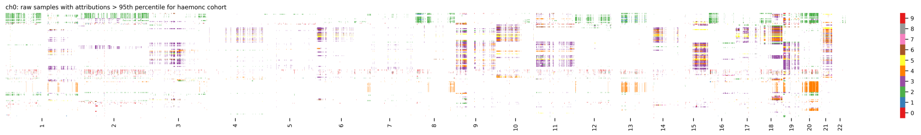


Supplementary Figure 64: Heatmap of CNA for samples in Haemonc cohort for total allele copy number. It highlights only high attribute loci as determined by the model, coloured according to CNA values. Loci not selected by the model are omitted, appearing as white spaces, to focus on regions of highest model attribution.

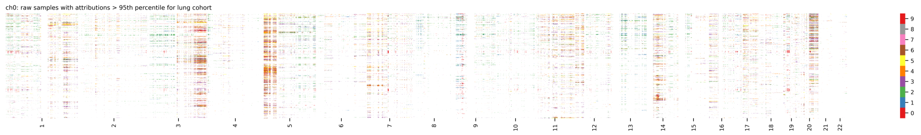


Supplementary Figure 65: Heatmap of CNA for samples in Lung cohort for total allele copy number. It highlights only high attribute loci as determined by the model, coloured according to CNA values. Loci not selected by the model are omitted, appearing as white spaces, to focus on regions of highest model attribution.
